# Spns1-dependent endocardial lysosomal function drives valve morphogenesis through Notch1-signaling

**DOI:** 10.1101/2024.03.26.586825

**Authors:** Myra N. Chávez, Prateek Arora, Alexander Ernst, Marco Meer, Rodrigo A. Morales, Nadia Mercader

## Abstract

Autophagy-lysosomal degradation is an evolutionarily conserved process key to cellular homeostasis, differentiation, and stress survival, which is particularly important for the cardiovascular system. Furthermore, experimental and clinical observations indicate it affects cardiac morphogenesis, including valve development. However, the cell-specificity and functional role of autophagic processes during heart development remain unclear. Here, we introduce novel zebrafish models to visualize autophagic vesicles *in vivo* and follow their temporal and cellular localization in the larval heart. We observed a significant accumulation of lysosomal vesicles in the developing atrioventricular and bulboventricular regions and their respective valves. Next, we addressed the role of lysosomal degradation using a Spinster homolog 1 (*spns1*) mutant. *spns1* mutants displayed morphological and functional cardiac defects, including abnormal endocardial organization, impaired valve formation and retrograde blood flow. Single-nuclear transcriptome analysis revealed endocardial-specific differences in the expression of lysosome-related genes and alterations of *notch1-*signalling in the mutant. Endocardial-specific overexpression of *spns1* and *notch1* rescued features of valve formation and function. Altogether, our study reveals a cell-autonomous role of lysosomal processing during cardiac valve formation upstream of *notch1-*signalling.

## Introduction

The macroautophagy/autophagy-lysosomal pathway plays a fundamental role in the maintenance of the cardiovascular system. Autophagy mediates the recycling of cytoplasmic components, organelle turnover, and the availability of basic building blocks and metabolites; it is active in all cells under basal conditions, but it can also be activated under stress conditions (Galluzzi *et al*, 2017; Klionsky *et al*, 2021b). During vertebrate embryonic development, autophagy-lysosomal processes are necessary to supply metabolites, enable organelle degradation for tissue remodelling and support cell differentiation (Boya *et al*, 2018; Perrotta *et al*, 2020; Allen & Baehrecke, 2020). In addition, autophagic processes have been suggested to control cell signalling driving tissue patterning and morphogenesis, with the lysosome acting as a cellular signalling centre (Lee *et al*, 2014; Allen & Baehrecke, 2020). Indeed, lysosomal function has been associated with congenital and acquired cardiovascular and valve diseases (Bhat & Li, 2021); and inherited lysosomal storage disorders have been linked to valve pathologies, such as pulmonary valve stenosis, mitral valve degeneration and valve calcification (Mueller *et al*, 2013; Ruiz-Guerrero & Barriales-Villa, 2018; Nordin *et al*, 2018; Platt *et al*, 2018). Moreover, links have been identified between autophagy-related gene expression and congenital atrioventricular valve defects (Yang *et al*, 2019; Lee *et al*, 2014). Yet, despite its evident biological and clinical significance, the function of lysosomal degradation during cardiac valve development has not been addressed.

Autophagic degradation requires the formation of an isolation organelle, known as the autophagosome, to sequester cytoplasmic components and target them to the lysosome, where the cargo is processed, and the resulting building blocks and degradation products are recycled or secreted. The lysosome contains digestive enzymes to break down cellular cargoes and is designed to maintain an acidic environment that allows their optimal activity (Bouhamdani *et al*, 2021). To study these highly dynamic processes during heart formation, experimental models that permit their precise *in vivo* observation are important assets. The zebrafish has long been considered an outstanding model organism for heart development studies thanks to its advantageous features for *in vivo* high-resolution imaging, such as external embryonic development, transparency, and amenability to genetic manipulation (González-Rosa, 2022). The similarities between its fast-developing heart and the mammalian heart make it particularly suitable to study the progression of congenital cardiovascular diseases. Also, many morphological characteristics of mammalian heart valves are shared with the zebrafish, and their formation is coordinated by the same signalling pathways that regulate mammalian endocardial cushion formation, endothelial-to-mesenchymal transformation, and valve interstitial cell proliferation (O’Donnell & Yutzey, 2020; Kemmler *et al*, 2021).

Moreover, the availability of cell- and organelle-specific zebrafish reporters and genome editing approaches have supported the study of processes underlying cardiovascular disease, including autophagy (Mathai *et al*, 2017; Zhang & Peterson, 2020; Klionsky *et al*, 2021a). The model has allowed to establish a link between autophagy-related gene expression and the correct transcriptional patterning of the early developing vertebrate heart (Lee *et al*, 2014), to mimic cardiac phenotypes observed in lysosomal storage disorders (Zhang & Peterson, 2020) and to functionally characterize mutations in lysosome-related genes associated with congenital cardiac defects (Ta-Shma *et al*, 2017; Lu *et al*, 2020). Still, the cell-type specific function of autophagy-lysosomal degradation during cardiac valve development is not fully understood and the developmental signalling pathways to which it is coupled remain to be elucidated.

Here we utilized high-resolution live imaging to describe in detail the autophagy-lysosomal events taking place during zebrafish heart development. We established a new knock-in autophagosome reporter and used cell- and organelle-specific transgenic lines to follow autophagosome and lysosome formation in the beating larval heart. Further, we described how autophagic degradation is connected to the cellular events taking place during cardiac valve formation.

Based on the *not really started* (*nrs*) zebrafish model, which carries a retroviral insertion affecting the expression of the gene *Spinster homolog 1* (*spns1*), we investigated how improper lysosomal function influences valve formation and function. *spns1* encodes a lipid-transporter that affects lysosomal acidity and, consequently, the function of pH-dependent lysosomal proteases (He *et al*, 2022; Scharenberg *et al*, 2023). The *nrs* mutant is a thoroughly characterized model of lysosomal-function impairment, which has allowed to study lysosomal deficiency in the contexts of embryogenesis (Young *et al*, 2002), senescence (Sasaki *et al*, 2014, 2017) and muscle degeneration (Coffey *et al*, 2021). Interestingly, the human *spns1* locus is located within the chromosome region 16p11.2, a region that has been associated with myxomatous mitral valve prolapse (Disse *et al*, 1999), aortic valve abnormalities (Ghebranious *et al*, 2007), septal defects and coarctation of aorta (Li *et al*, 2017). Using light-sheet, confocal and electron microscopy, we compared the morphology and function of mutant and sibling hearts and found that lysosome activity in endocardial cells is required for proper valve morphogenesis. Then, based on single-nuclei transcriptome analysis of the *nrs* mutant heart, we exposed a role for lysosomal function in myocardial-endocardial cell interaction mediated through the *notch1* signalling and propose a link between the lysosomal function and correct valve development.

## Results

### Autophagosome formation accompanies valve development

To elucidate the role of autophagy-lysosomal processing during cardiac valve formation, we characterized the kinetics of autophagosome and lysosome accumulation in the developing zebrafish heart (Fig. 1A and B). For this, we established and made use of zebrafish transgenic models and live fluorescent dyes to locate autophagosome and lysosome vesicles within different tissues and structures of the larval heart. Furthermore, we optimized a previously described (Marques *et al*, 2022) experimental setup for *in vivo* imaging of zebrafish larvae along with a downstream image processing workflow to get a three-dimensional reconstruction of the imaged beating larval heart (Fig. 1C). This allowed us to identify the nascent valves in the outflow tract (OFT) and atrioventricular canal (AVC), which is only accurately possible when their movement can be observed (Vignes *et al*, 2022). We performed live imaging at four larval stages, during which the main cellular processes that lead to valve formation take place (Steed *et al*, 2016b; Gunawan *et al*, 2020): endocardial cell (EnC) clustering at the AVC (48 hours post-fertilization, hpf), EnC migration into the cardiac jelly (56 hpf), formation of a multilayered primitive valvular structure (72 hpf) and elongation of the valve leaflet (96 hpf).

**Figure 1.**
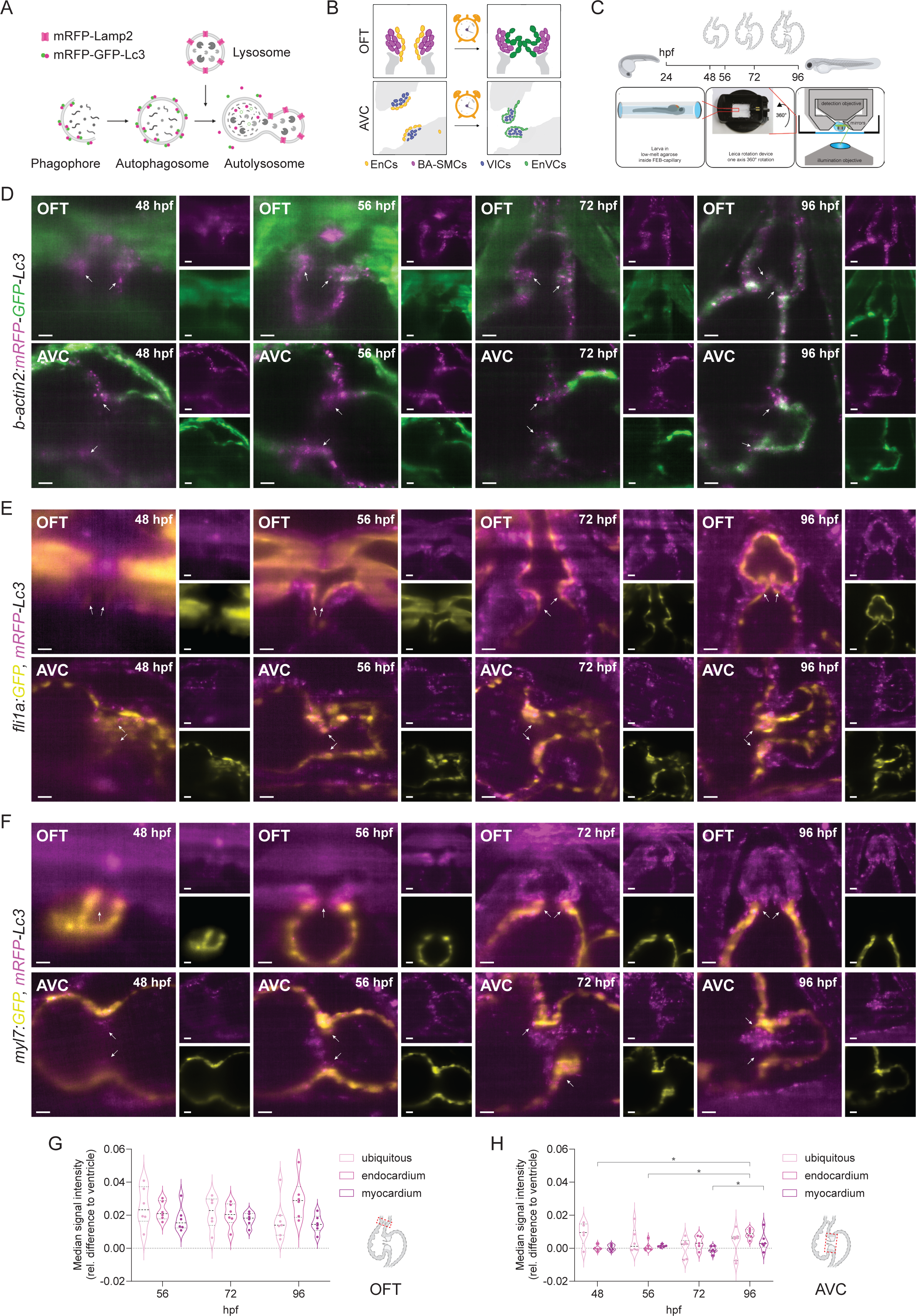
Autophagosome formation accompanies valve development. (**A**) Schematic overview of the process of autophagy-lysosomal degradation involving phagosome and autophagosome formation, fusion with lysosome, and lysosomal degradation. The transgenic autophagosome and lysosome fluorescently tagged reporter proteins used in this study are illustrated. (**B**) Graphic representation of the central question of the study: when and where are autophagy and lysosomal processes taking place during cardiac valve development (OFT: outflow tract, AVC: atrioventricular canal, EnCs: endocardial cells, BA-SMCs: Bulbus arteriosus (BA) smooth muscle cells, VIC: valve interstitial cells, EnVCs: endocardial valve cells) (**C**) Experimental plan followed for *in vivo* imaging of autophagic vesicles in the developing zebrafish heart with an overview of the imaging setup. Transgenic zebrafish larvae were imaged at 48, 56, 72 and 96 hpf upon 3 h treatment with chloroquine (2 mM) to observe autophagosomes and lysosomes in the beating heart using light-sheet microscopy and following a 3D-t acquisition mode. Image reconstruction allows to observe the distribution of autophagic vesicles in the larval heart at different developmental stages (**D-F**) Reconstructed image acquisitions by 3D-t light-sheet microscopy. Stack projections (**D**) and optical sections through hearts (**E and F**) at the indicated developmental stages are shown. Anterior is to the top (V: ventricle, A: atrium) (**D**) *Tg(b-actin2:mRFP-GFP-Lc3)* zebrafish larvae show mRFP^+^-autophagosomes/autolysosomes ubiquitously distributed in the developing heart and the accumulation of mRFP-GFP^+^ autophagosomes in the atrioventricular canal (AVC) and outflow tract (OFT) between 72 and 96 hpf respectively. White arrows point to the developing bulboventricular (upper inset) and atrioventricular (bottom inset) valves. (**E and F**) The new transgenic knock-in line *Tg(mRFP-Lc3)* expresses an mRFP-tagged version of the endogenous autophagosome backbone protein Lc3. This reporter line allows the unbiased observation of mRFP^+^-autophagosome formation and their localization in endocardial (*Tg(fli1a:GFP)*) and myocardial (*Tg(myl7:GFP)*) tissues, and revealed their accumulation in the OFT and AVC regions of the developing heart at 72-96 hpf (white arrows, V: ventricle) (**G and H**) Quantification of mRFP^+^-median fluorescence in endocardial and myocardial tissues in the OFT and AVC compared to the ventricle (rel. difference to ventricle > 0). Scale bars represent 12.5 µm in insets and 25 µm in all other figures. Each point represents one larva (Kruskal-Wallis test, *p ≤ 0.05).

Using the ubiquitous dual-fluorophore transgenic line *Tg(b-actin2:mRFP-GFP-Lc3)* (Chávez *et al*, 2020) that expresses the autophagosome microtubule-associated protein light chain 3B (MAP1LC3B, hereafter referred as Lc3) tagged with both a monomeric red (mRFP) and green fluorescent protein (GFP), we first followed the distribution of autophagosomes (mRFP-GFP^+^) and autolysosomes (mRFP^+^) in the larval heart (Fig. 1D, Video S1). Prior to imaging, larvae were shortly treated with the lysosomal-inhibitor chloroquine to block autophagic flux (Mauthe *et al*, 2018) and allow the observation of autophagosome puncta, following similar protocols used to observe autophagy activation in zebrafish models (Mathai *et al*, 2017; Chávez *et al*, 2020). The cytoplasmatic signal of GFP-Lc3 was uniformly observed throughout the heart, though very few GFP-puncta were discernible (Fig. 1D). In contrast, mRFP-labelled autophagosomes/autolysosomes were detected as discrete puncta across the heart from 48 hpf onwards. We therefore decided to consider the median fluorescence signal of mRFP as the most robust parameter to evaluate relative regional differences in autophagosome accumulation during cardiac development.

A caveat with the use of a ubiquitous line such as *Tg(b-actin2:mRFP-GFP-Lc3)* is that tissues with particularly high *b-actin2* promoter activity, such as the myocardium, will show higher reporter expression as compared to the less-contractile endocardium. To overcome this limitation and accomplish an accurate tracking of autophagosome accumulation in all cardiac tissues, we decided to create a knock-in transgenic line (*TgKI(mRFP-map1lc3b)*, hereafter referred as *TgKI(mRFP-Lc3*)) that allows the visualization of the endogenous Lc3, and as a result, the unbiased tracking of autophagosome vesicles (Fig. S1A-D). For this, we introduced the coding sequence of the fluorescent protein mRFP upstream of and in frame with the first exon of the *map1lc3b* zebrafish orthologue, to generate a tagged native protein (Wu *et al*, 2018). By crossing this new line with endocardial (*Tg(fli1a:GFP)*, Fig. 1E, Video S2) and myocardial (*Tg(myl7:GFP)*, Fig. 1F, Video S3) reporters, we could locate autophagosomes in the myocardium and endocardium.

Both qualitative observations (Fig. 1D-F) and quantitative mRFP-fluorescence estimations (Fig. S1E) indicated an overall increase in autophagosome formation/accumulation during heart development. However, compared to the rest of the ventricle, the mRFP-signal in the OFT was consistently higher throughout the development of the heart (Fig. 1G, rel. difference > 0). Also, in this region, mRFP-Lc3 expression was not restricted to the myocardium and endocardium but was also detected in periostin-positive cells (*postnb^+^*) lining the OFT surface from 72 hpf onwards (Fig. S1F), thus suggesting an overall increased autophagic activity in this heart structure. In the AVC, we identified a rise in both ubiquitous and endogenous mRFP-Lc3 signal starting from 72 hpf (Fig. 1E and F white arrows), which became significant both in the atrioventricular myocardium and endocardium by 96 hpf when compared to earlier developmental stages (Fig. 1H).

### Lysosome accumulation during valve development

To follow the complete autophagic pathway, in a next step we tracked lysosome accumulation using LysoTracker™ Deep Red (Fig. 2A, Video S4), which allows to fluorescently label acidic vesicles *in vivo*. To observe their co-localization with autophagosomes, we used the ubiquitous autophagosome reporter *Tg(cmv:EGFP-Lc3)*. Also, because the LysoTracker™ dye may also label acidic endosomes and its fluorescence depends on the dye permeability and environmental pH (Klionsky *et al*, 2021a), we complemented our characterization with a transgenic Lamp2-reporter *TgBAC (lamp2:RFP)* to identify lysosomes in endocardial cells (*Tg(kdlr:EGFP-CAAX)*, Fig. 2B, Video S5) and in cardiomyocytes (*Tg(myl7:GFP)*, Fig. 2C, Video S6). We considered the median fluorescence signal of LysoTracker™ Deep Red and mRFP to quantify lysosomal vesicles in the developing heart (Fig. S1F) and evaluate relative regional differences in lysosome accumulation (Fig. 2D, E).

**Figure 2:**
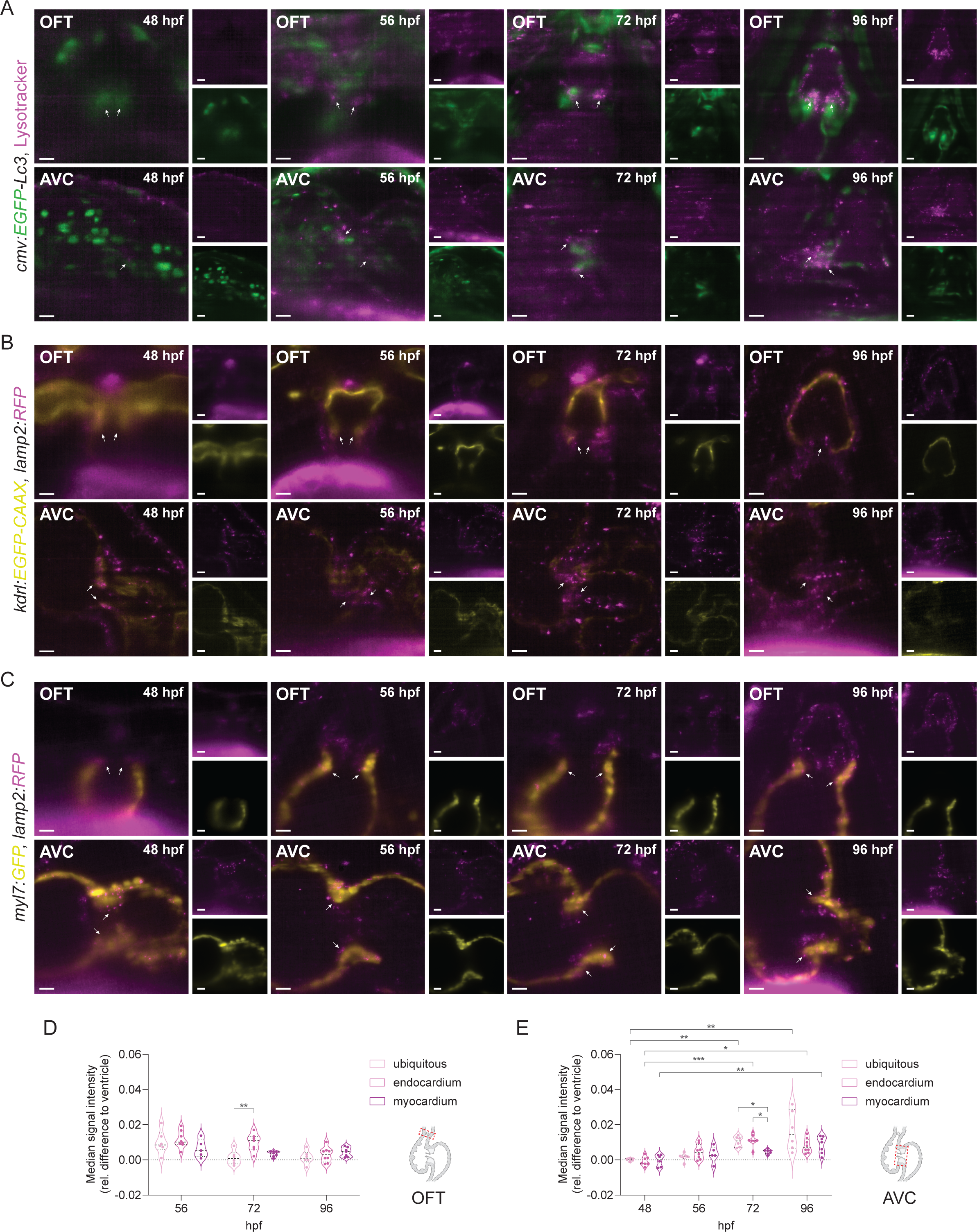
Lysosome accumulation during heart valve development. (**A-C**) Reconstructed live image acquisitions by 3D-t light-sheet microscopy. Stack projections (**A**) and optical sections through hearts (**B and C**) at the indicated developmental stages. Anterior is to the top (V: ventricle, A: atrium). (**A**) *Tg(cmv:EGFP-Lc3)* embryos were stained with LysoTracker^TM^ Deep Red to observe GFP^+^ autophagosome and LysoTracker^+^ acidic lysosomal vesicles in the developing heart. White arrows point to the developing bulboventricular (upper inset) and atrioventricular (bottom inset) valves. EGFP signal is strongly detected in circulating erythrocytes between 48 and 56 hpf, but later localizes mainly in the OFT and AVC. LysoTracker-labelled lysosomes are scarcely observed before 56 hpf in cardiac tissues, but begin to accumulate in the OFT and AVC at 72 hpf and are prominent in the developing valves at 96 hpf. (**B and C**) The lysosome reporter *Tg(lamp2:RFP)* revealed the presence of lysosomal vesicles in 48-56 hpf hearts in both the OFT and AVC (white arrows) starting from 72 hpf in endocardial cells (*Tg(fli1a:GFP)*) and at 96 hpf in cardiomyocytes (*Tg(myl7:GFP)*) (**D and E**) Quantification of the median fluorescence signal of the LysoTracker^+^ and mRFP^+^-lysosomes exposed a significant accumulation of lysosomes in endocardial cells in the OFT (**D**) at 72 hpf, and overall higher lysosomal activity in all cardiac tissues of the AVC from 72 hpf onwards. Scale bars represent 12.5 µm in insets and 25 µm in all other figures. Each point represents one larva (Kruskal-Wallis test, *p ≤ 0.05, **p ≤ 0.01, ***p ≤ 0.001).

Contrary to the *b-actin2*-mediated mRFP-GFP autophagosome reporter, the *cmv*-transgenic line showed high expression in erythrocytes at 48 and 56 hpf. Nonetheless, from 72 hpf onwards, cells in the OFT and AVC regions showed high green fluorescence and some distinguishable EGFP-Lc3^+^ puncta, confirming our observations of autophagy activation with the ubiquitous and the endogenous LC3-reporters in these regions (Fig. 1). LysoTracker™-vesicles, on the other hand, were sparsely found in the ventricle at 48 hpf, yet starting from 56 hpf, a significant gradual increase in LysoTracker™-labelled puncta was observed throughout the larval heart (Fig. 2A, Fig. S1G). Co-localization of EGFP^+^ and LysoTracker™ puncta could be observed both in the OFT and AVC nascent valves from 72 hpf onwards (Fig. 2A, white pixels), indicating autophagosome lysosomal processing during primitive valve formation and valve elongation. Also, we noticed that both newly formed mRFP-GFP^+^ autophagosomes (Fig. 1D, Video S1, white puncta) and LysoTracker™-colocalizing EGFP-Lc3^+^ autophagosomes (Fig. 2A, Video S4, white puncta) were enriched at the base of the OFT and AVC valves at 96 hpf, suggesting a high degree of autophagy-mediated cellular remodelling (Allen & Baehrecke, 2020).

In contrast to LysoTracker™-stained lysosomal vesicles (Fig. 2A), *lamp2*^+^-lysosomes could already be observed at 48 hpf across the larval heart (Fig. 2B,C) and their numbers appeared to be comparable among developmental stages (Fig. S1F). Yet, when focusing on the atrioventricular region, we found a significant accumulation of *lamp2*^+^-lysosomes (Fig. 2B, C, E) in both endocardial (from 72 hpf onwards) and myocardial cells (at 96 hpf). This was consistent with the significant accumulation of LysoTracker™ vesicles observed at 72 and 96 hpf (Fig. 2A, E). We also observed a significant accumulation of *lamp2*^+^-lysosomes in OFT endocardial cells at 72 hpf (Fig. 2B, D), and co-localization of LysoTracker™-puncta with neighbouring *postnb*^+^-cells (Fig. S1H) in this region at this stage. Altogether, these results point towards a sudden localized rise in both autophagosome and lysosome accumulation during the formation and elongation of both the OFT and AVC valves (72-96 hpf).

### Impaired lysosomal degradation in the developing *nrs* mutant heart

To understand the function of autophagy-lysosome degradation in the development of the heart, we used the *nrs* zebrafish mutant (Young *et al*, 2002), which is a thoroughly characterized model of lysosomal-function impairment (Sasaki *et al*, 2017). Following our established experimental workflow to observe autophagic vesicles in the developing heart, we evaluated autophagic processing in the *nrs* mutant compared to homozygous/heterozygous siblings (Fig. 3A-C). Consistent with previous observations in the *nrs* yolk (Young *et al*, 2002; Sasaki *et al*, 2014) and skeletal muscle (Coffey *et al*, 2021), we found a significant accumulation of mRFP^+^-autolysosomes, LysoTracker™-vesicles and lamp2^+^-lysosomes in the mutant heart at 3 days post-fertilization (dpf, Fig. 3D), where both autophagosomes/autolysosomes and lysosomes were particularly enriched and enlarged in the OFT and AVC regions compared to siblings (Fig. 3D, white arrows). These observations are consistent with the enlarged non-degraded autolysosomal compartments previously reported in the *spns1* mutant, which have been attributed to impaired autolysosomal acidification (Sasaki *et al*, 2014).

**Figure 3.**
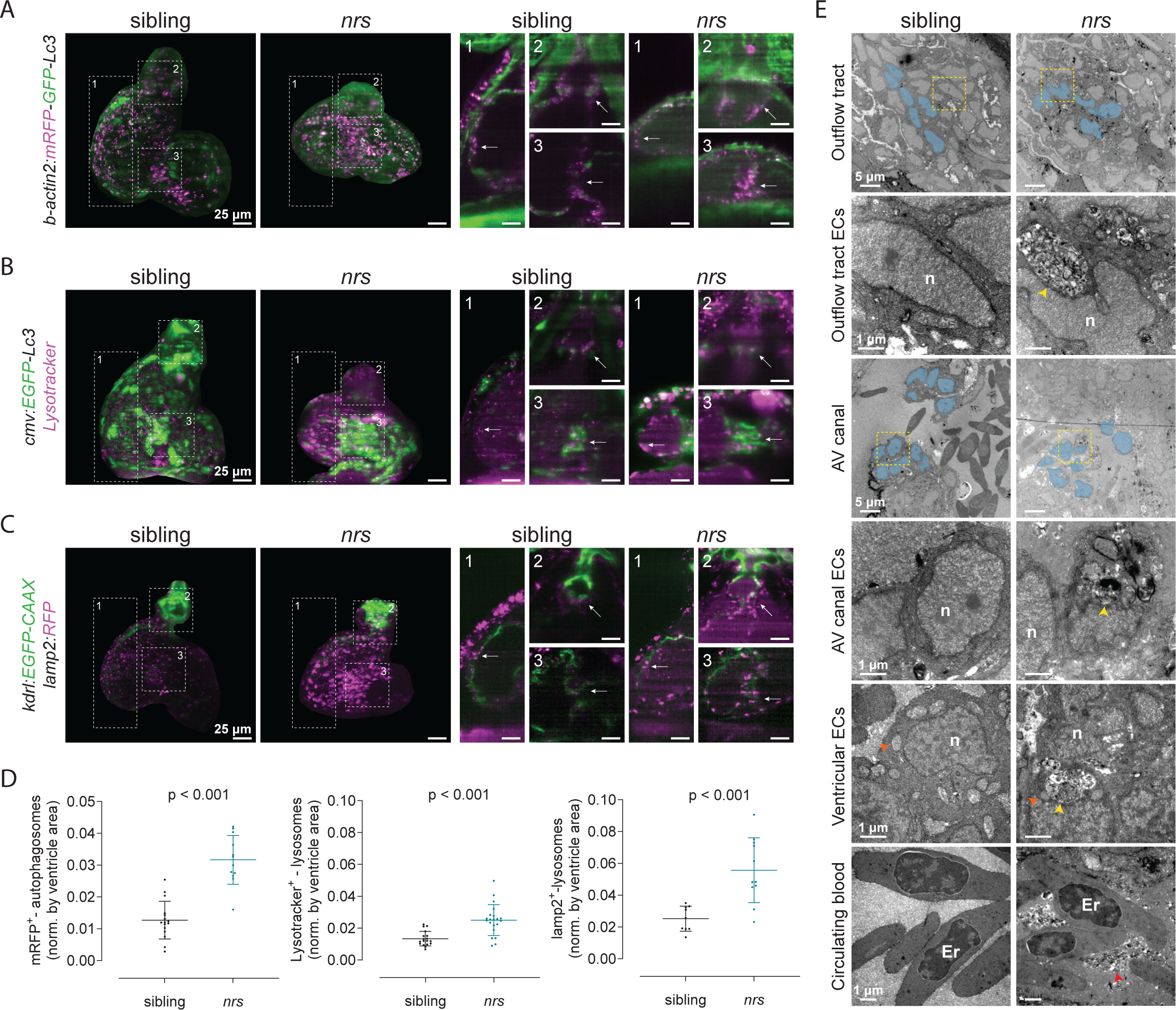
Impaired lysosomal degradation in the developing *nrs*-mutant heart. (**A-C**) The transgenic lines *Tg(b-actin2:mRFP-GFP-Lc3)*, *Tg(cmv:EGFP-Lc3)* and *Tg(kdlr:EGFP-CAAX; lamp2:RFP)* were crossed into the *nrs* mutant background to assess autophagic processing in the larval heart at 3 dpf by light-sheet microscopy. Projections and optical sections of reconstructed 3D-t acquisitions are shown. (**D**) The number of puncta was estimated for (**A**) mRFP^+^-autophagosomes/autolysosomes, (**B**) LysoTracker^+^ lysosomes and (**C**) lamp2^+^ lysosomes and normalized to the ventricle cross-sectional area. The results show a significant accumulation of all autophagic vesicles in *nrs*-mutant hearts compared to their siblings, which indicates an impairment in autophagic flux. Above all, mRFP^+^ autophagosomes/autolysosomes were densely distributed within the AVC (**A**, white arrows), whereas lysosomes were enriched and enlarged in both the mutant OFT and the AVC (**B and C**, white arrows). Each point represents one larva. Statistical analysis was performed using Welch’s t-test. Scale bars represent 25 µm. (**E**) Transmission electron microscopy of 4 dpf mutant and sibling hearts revealed the presence of massive lysosomal compartments with partially degraded contents in *nrs* hearts (yellow arrowheads), mostly in endothelial cells in the OFT and AVC (N=3). In addition, compared to sibling ventricular and atrioventricular endocardial valve cells (blue-coloured nuclei), which show high organization and homogenous shape mutant endocardial cells were amorphous and less organized. Also, in sibling endocardial cells, mitochondria were located around the nucleus (orange arrowheads), whereas they could hardly be distinguished inside mutant endocardial cells. Lysosomal-derived contents were observed in the circulating blood in the *nrs* hearts lumen (red arrowheads).

To confirm our observations, larval samples were processed for transmission electron microscopy and ultrathin sections containing OFT and AVC regions were obtained from sibling and mutant hearts (Fig. 3E). Mutant hearts revealed a disorganization of the OFT and AVC tissues and high accumulation of intracellular vesicles that are not present in control siblings. At subcellular resolution, we found massive lysosomal vesicles containing partially degraded cargoes above all in endothelial cells of the OFT and AVC (yellow arrowheads). Furthermore, the visible organization of endothelial cells delineating the developing valves in the sibling heart, was missing in the mutant heart, and the respective elongated and square shape of the sibling OFT and AVC endothelial valve cells was lost (Fig. 3E, cells with blue-coloured nuclei). Also, while ventricular endocardial cells showed a homogenous distribution of mitochondria surrounding their nucleus (orange arrowheads), mitochondria were scarcely found in mutant endocardial cells, and then mainly near lysosomal vesicles. We did not observe changes in the appearance of cardiomyocyte mitochondria or actin fibres in *nrs* hearts (Fig. S2A), though we noted that lysosome and autophagosome vesicles were more abundant in cardiomyocytes close to the OFT and AVC in both sibling and mutant hearts. Even though massive lysosomal vesicles were rare in mutant cardiomyocytes compared to endocardial cells, we observed dark structures full of double membrane autophagic vacuoles within the mutant myocardium suggesting accumulation of unprocessed lysosomal contents (yellow arrowheads). In addition, we observed vesicle-like particles in the mutant heart lumen with high similarity to the lysosomal contents accumulated in endocardial cells (Fig. 3E, red arrowheads). In line with the described role of *spns1* in lysosomal function (Sasaki *et al*, 2014; He *et al*, 2022; Scharenberg *et al*, 2023), our results point towards a significant impairment in lysosomal processing also in *nrs* hearts, particularly affecting endocardial tissues and the organization of endothelial valve cells.

### Lysosomal mutants present abnormal valve development and function

To assess the impact of lysosomal dysfunction in the developing larval heart, we compared the cardiac morphology and function of *nrs* mutant to their siblings at 3 and 4 dpf (Fig. 4, Fig. S2B-G). We observed a significantly reduced ventricle size and an overall rounder ventricular shape at 3 dpf in the mutant heart, which did not recover by 4 dpf (Fig. S2B and C). The length of the OFT was proportionally smaller in the mutant heart at 3 dpf, though no differences were found at 4 dpf (Fig. 4A and B). In contrast, the AVC was proportionally wider in the mutant at both developmental time points (Fig. 4A and B). The mutant endocardium appeared to collapse inside the ventricle lumen at 4 dpf, even though it showed a comparable organization to the sibling at 3 dpf (Fig. 4 and B, Video S7). Moreover, while myocardial contractility appeared similar between siblings and *nrs* at 4 dpf, in the mutant heart the movement of the endocardial tissue was restrained, and obstruction was observed in the outflow tract (Video S7).

**Figure 4.**
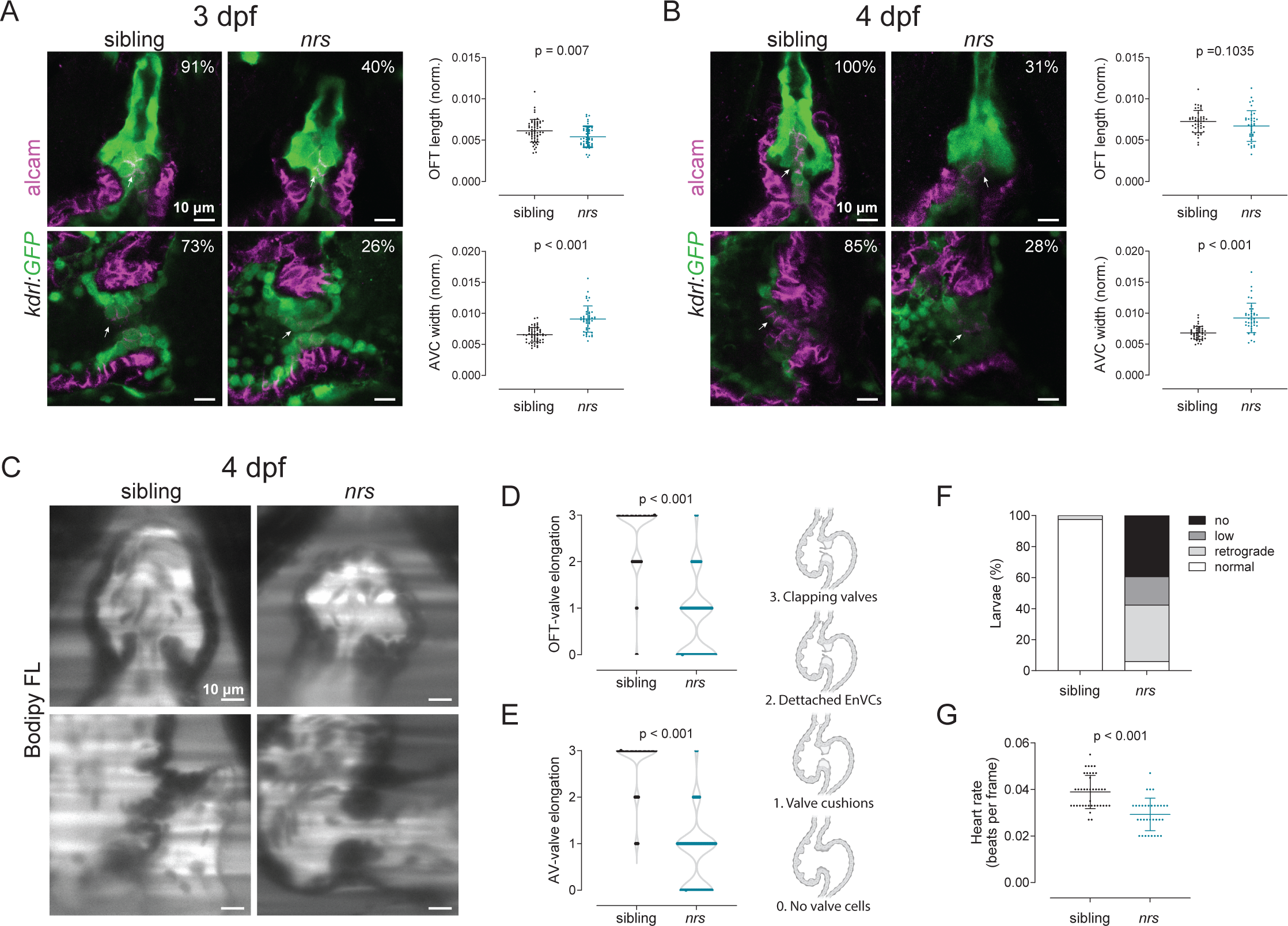
*nrs* mutant have abnormal valve development and function. (**A and B**) Cardiac morphology and valve development was evaluated in *Tg(kdrl:GFP)* transgenic *nrs* mutants between 3 and 4 dpf. Confocal microscopy images of whole mount immunostained larval hearts are shown. When normalized to the ventricle cross-sectional area, the length of the OFT in the mutant heart was proportionally smaller at 3 dpf in *nrs* larvae, whereas the AVC was significantly wider compared to their siblings at both experimental time-points. Most sibling hearts displayed alcam^+^-organized primitive valve layers at 3 dpf (OFT 91%, AVC 73%) and aligned alcam^+^-valve endothelial cells at 4 dpf (OFT 100%, AVC 85%). In contrast, alcam^+^-endocardial cells were rarely observed in mutant larvae both at 3 dpf (OFT 40%, AVC 26%) and 4 dpf (OFT 31%, AVC 28%) and the organization of valve endothelial cells was lost by 4 dpf. Each dot represents one larva. Statistical analysis was performed using Mann-Whitney-U test. (**C**) To evaluate valve function and blood flow, sibling and mutant larvae were immersed in BODIPY™ FL C5-Ceramide, a green-fluorescent dye which counterstains the blood plasma and live imaged by light-sheet microscopy. Qualitative parameters were considered to evaluate (**D**) OFT-and (**E**) AV-valve elongation and function (1: valve cushions, 2: delaminated valve cells, 3: elongated clapping valves. Each dot represents one valve. Chi-square test was applied to analyse differences between experimental groups. (**F**) Heart rate was calculated from the number of beats during the acquisition of 150 frames. Each dot represents one larva. Statistical analysis was performed using Mann-Whitney-U test. (**G**) Percentage of larvae presenting normal, low, retrograde or no blood flow.

Both valve morphology and function were severely affected in mutant hearts. While endocardial *alcam^+^*-cell, layers which give rise to the OFT valves, were present in almost all the siblings, they were found in less than 40% of the mutants at 3 dpf and in only ∼30% at 4 dpf (Fig. 4A and B). In the case of the AV-valves, which consist of aligned cubic alcam^+^-endocardial cells both at 3 and 4 dpf, only ∼26-28% of the larvae showed alcam expression in atrioventricular endocardial valve cells (EnVCs), and their organization was lost (Fig. 4A and B). Functional elongated and clapping valves were rarely found in the mutant heart, and correspondingly, a high prevalence of retrograde flow was often observed (Fig. 4C-F and Video S8). In addition, lower heart rate was observed in the mutant hearts both at 3 (Fig. S2D) and 4 dpf (Fig. 4G), and a reduced cardiac output (Fig. S2E), consistent with the decreased *nrs* ventricle size estimated at 3 dpf. Yet, no evidence for ejection fraction shortening (Fig. S2F) or arrythmia (RMSSD, Fig. S2G) was found.

To assess the cardiac susceptibility to mutations in *spns1*, we evaluated cardiac morphology and function in *spns1*-crispants (Fig. S3). Compared to mutants, *spns1* crispants had an ameliorated of yolk opacity and a higher degree of yolk extension (Fig. S3A), which have been correlated with a decreased severity of the mutant phenotype (Sasaki *et al*, 2017). Despite this, *spns1* crispants still showed a significant reduction in ventricle size and an increase in circularity index compared to control injected larvae (Fig. S3B-D). Like the mutants, the formation of organized valve leaflets was affected in the crispants, with only 36% of the larvae showing alcam^+^-endothelial cells delineating the nascent atrioventricular valve leaflets, compared to 68% of the control larvae (Fig. S3E). As in the mutant, both OFT and AV valve elongation were diminished by *spns1* deficiency (Fig. S3F-H) and thickened non-clapping valves were often found in the crispants, which partially affected unidirectional blood flow (Fig. S3I). However, in contrast to the mutant, *spns1* crispants showed no difference in heart rate compared to controls, which would indicate that impaired valve development in *spns1*-deficient larvae is independent of ventricular function (Fig. S3J). Altogether our results suggest that loss of *spns1* function compromises cardiac development, particularly affecting valve morphogenesis and function.

### Endocardial specific overexpression of *spns1* rescues valve development and function

Since our results suggest that lysosomal impairment particularly affects endocardial development and function, and that the absence of *spns1* particularly disrupts lysosomal processes in the endocardial tissue, we investigated whether the endocardial-specific expression of the wild type *spns1* gene could rescue the observed cardiac phenotypes. For this, we established a UAS-mediated transgenic line that permitted *spns1* expression when crossed with a fli1a-Gal4 line (Fig. 5). As above, we obtained double transgenic mutant larvae by crossing single transgenic heterozygous parents (Fig. 5A., *spns1^+/-^, fli1a:Gal4* and *spns1^+/-^*, *UAS:spns1*). When we verified the expression of *spns1* in the mutants via qPCR (Fig. 5B), we found that most of the living *nrs*-mutants at 4 dpf were double transgenic *(fli1a:Gal4*, *UAS:spns1*), hinting towards a rescue effect in *nrs* survival (Fig. 5C). While endocardial overexpression of *spns1* did not induce any significant morphological or functional change in the ventricle (Fig. 5D), it led to a partial rescue of valve elongation in the mutant valves (Fig. 5E-G). This resulted in valve function recovery, as shown by an increased percentage of larvae with normal blood flow (Fig. 5H, Video S9), which could not be attributed to heart function recovery (Fig. 5I). Moreover, electron microscopy revealed a reduction in the frequency and size of lysosomal vesicles with partially degraded cargo in endocardial cells (Fig. 5J, K). Both endocardial appearance and organization, as well as the perinuclear distribution of their mitochondria were improved (Fig. 5J, orange arrowheads). Even though lysosomal contents were still found in the cardiac lumen, the endocardial overexpression of *spns1* reduced partially degraded lysosomal contents within the myocardium (Fig. 5L). In summary, the results of this study suggest that *spns1*-dependent lysosomal function is required in endocardial cells acting upstream of *notch1* signalling to regulate correct myocardial-endocardial cell interaction and organization and drive valve morphogenesis.

**Figure 5.**
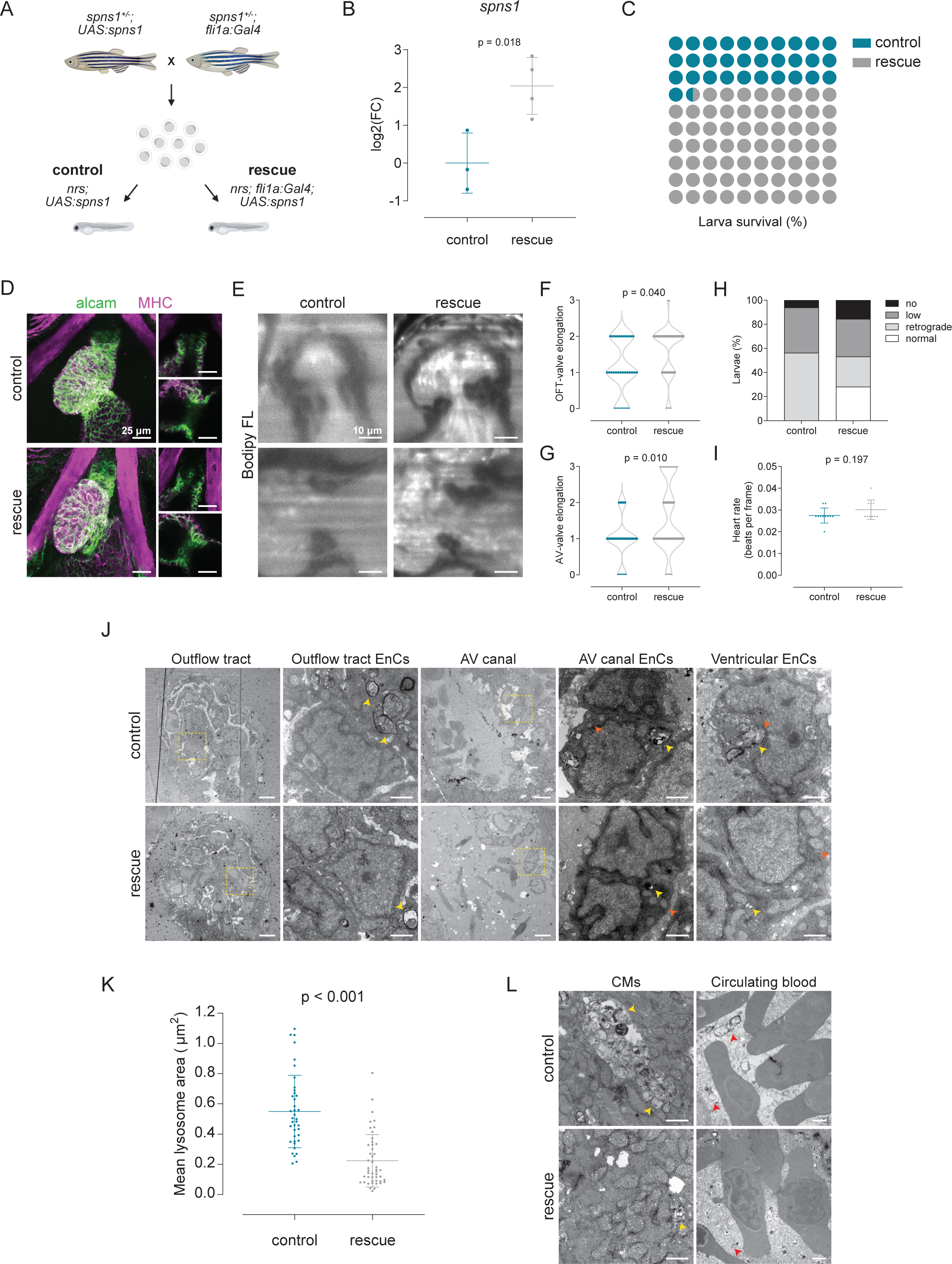
Endothelial specific overexpression of *spns1* rescues valve development and function. (**A**) To address the endocardial-specific susceptibility towards lysosomal impairment, a rescue strategy based on the endothelial-specific overexpression of the wild-type *spns1* gene was implemented. For this, single transgenic heterozygous parents (*spns1^+/-^;fli1a:Gal4* and *spns1^+/-^;UAS:spns1*) were crossed to obtain control (*nrs;UAS:spns1*) and rescue (*nrs;fli1a:Gal4 UAS:spns1*) larvae. (**B**) Overexpression of the *spns1* was verified by qPCR. (**C**) Larval survival percentage at 4 dpf for each experimental group. (**D**) Confocal microscopy images of wholemount immunostaining of larval hearts show enhanced alcam expression in the OFT and AVC in rescued hearts. Scale bars represent 25 µm. (**E**) Light-sheet microscopy of BODIPY™ FL C5-stained larvae was performed to evaluate cardiac and valve function. (**F**) OFT-and (**G**) AV-valve elongation and function were evaluated qualitatively (1: valve cushions, 2: delaminated valve cells, 3: elongated clapping valves. Each dot represents one valve. Chi-square test was applied to analyse differences between experimental groups. (**H**) Percentage of larvae presenting normal, low, retrograde or no blood flow. (**I**) Heart rate was calculated from the number of beats during the acquisition of 150 frames. Each dot represents one larva. Statistical analysis was performed using Mann-Whitney-U test. (**J**, **K**) Transmission electron microscopy revealed a reduction in the accumulation of partially degraded lysosomal contents in endocardial cells of rescued larvae (yellow arrowheads), and a partial restoration of endocardial morphology. Also, the alignment and shape of endocardial cells and the preservation of their mitochondria (orange arrowheads) was reinstated through the overexpression of *spns1*. (**I**) Accumulation of lysosomal contents was reduced in myocardial cells (CMs, yellow arrowheads, yet lysosomal contents in the blood plasma were still present in the rescue group (red arrowheads). N ≥ 3 per experimental group. (**M**) Graphical representation of the location and developmental stage in which autophagy-lysosomal processes take place during cardiac valve development and the role that Spns1 plays in preserving lysosomal function.

### Expression of lysosome related genes and *notch1b*-signalling is affected in *nrs*-mutants

To understand the transcriptomic programs affected by lysosomal dysfunction in *nrs* hearts, we performed a comparative analysis between sibling and mutant hearts at 3 dpf using single-nuclei RNAseq, (snRNAseq). We identified 18 different cell populations including 3 myocardial and 5 endocardial subclusters (Fig. 6A, Fig. S4A-C, Table S1). To identify the endocardial valve cells (EnVCs) among the latter, we compared the expression of annotated genes involved in valve development or genes which have been previously suggested as valve cell markers (Grimes & Kirby, 2009; Chen *et al*, 2013; Kuleshov *et al*, 2016; Goddard *et al*, 2017; Burkhard & Bakkers, 2018; Duchemin *et al*, 2019; Fontana *et al*, 2020; Xie *et al*, 2021; Ma *et al*, 2021; Queen *et al*, 2023) (Table S2), namely *notch1b*, *nrg1*, *rspo2*, *alcama*, *cdh5* and *has2* (EnVCs, Fig. S2B). Similarly, based on the expression of atrioventricular cell markers (*bmpr2b*, *col1a1a*, *robo1*, *hapln1b*, *notch1b*, *ednraa*, *col1a2*, *fn1a*), we identified one myocardial subcluster as atrioventricular cardiomyocytes (CMs AVC, Fig. S2C). When comparing the cell populations between siblings and mutants, we observed that, in contrast to other cell types, the proportion of myocardial and endocardial cells was reduced in the subclusters that had highest valve development-related gene expression (CMs AVC, EnCs 3, EnCs 4 and EnVCs), as well as neurons (Fig. 6B), suggesting higher susceptibility to *spns1* impairment in these populations.

**Figure 6.**
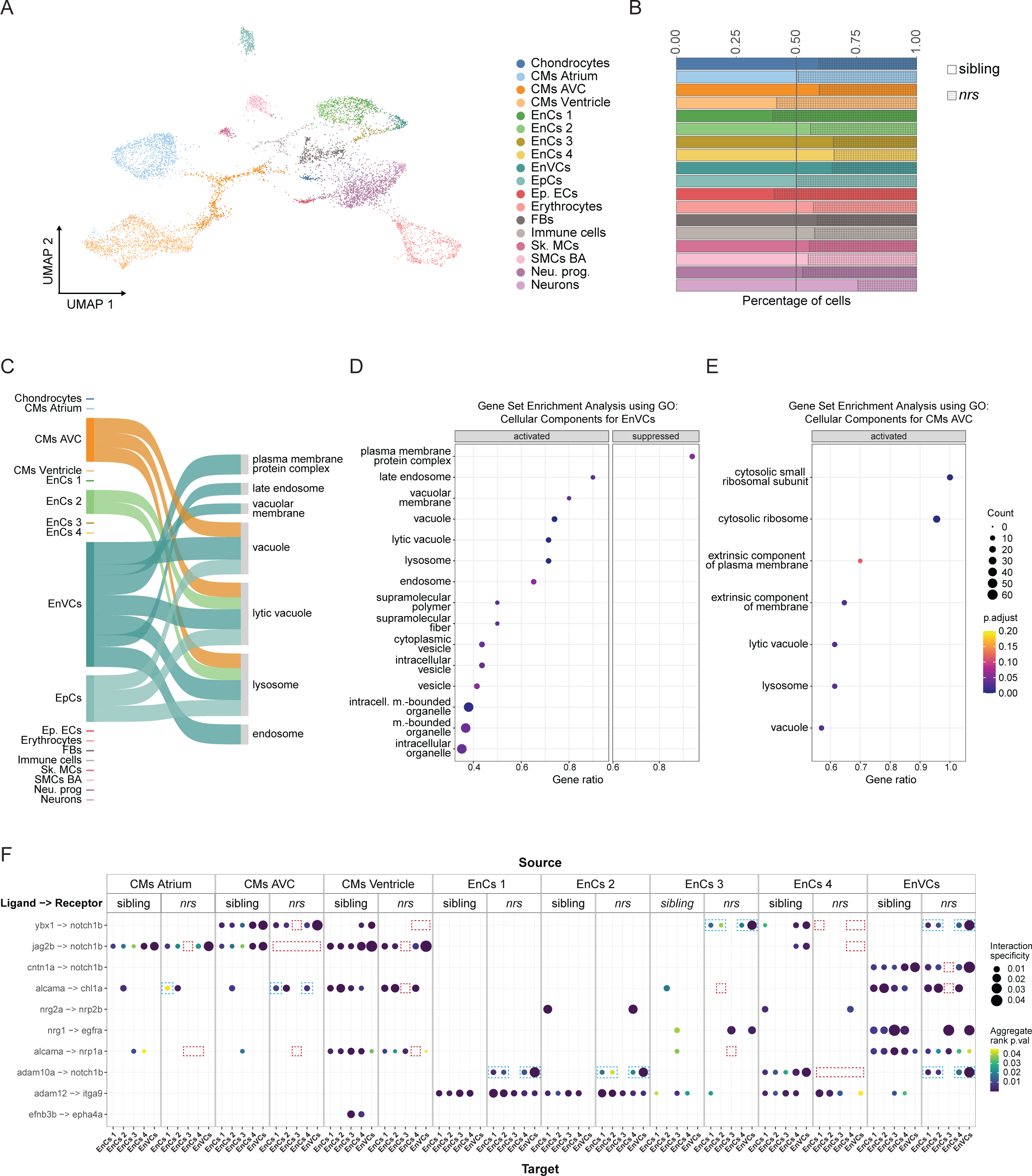
Expression of lysosome related genes and notch1b-signalling is affected in *nrs*-mutants. Results obtained from a snRNA transcriptome analysis of sibling and *nrs* mutant hearts at 3 dpf. (**A**) UMAP showing the various identified cell types from the snRNASeq performed for mutant and sibling hearts at 3 dpf. (**B**) The proportion of different cell types observed in sibling versus mutant suggest that lysosomal impairment above all affects the representation of atrioventricular cardiomyocytes (CMs AVC), three endocardial populations (EnCs 3, EnCs 4) among them endocardial valve cells (EnVCs) and neurons in the larval heart. (**C**). Sankey diagram illustrates the cell-specificity of overrepresented lysosomal and membrane related gene pathways found in *nrs* mutants compared to siblings highlighting the relevance of lysosomal degradation in EnVCs. The width of the arrows is proportional to the number of differentially expressed genes in each cell cluster. (**D and E**) Gene set Enrichment Analysis using the differentially expressed genes between siblings versus mutants in EnVCs and (**E**) atrioventricular CMs, where ‘activated’ and ‘suppressed’ represent the pathways that are activated or suppressed in the mutant condition. The results obtained suggest an upregulated transcriptional response in mutant EnVCs and atrioventricular CMs to compensate impaired lysosomal function. (**F)** Ligand-receptor analysis from LIANA depicting the ligands from the source cells (top) to target cells (bottom) with ligand and receiving receptor shown on the y-axis in the sibling and the mutant conditions. The ligand-receptor interactions affecting *notch1b*-and *alcama*-signalling are highlighted for the *nrs* mutant (red square = missing interaction, blue square = differing interaction).

To gain a deeper understanding of the pathways affected in the mutant heart, we performed a gene set enrichment analysis (GSEA) on the differentially expressed genes of each cell type (Fig. 6C-E, Table S4). GSEA uses a ranked list of differentially expressed genes to calculate an enrichment score for each pathway (Subramanian *et al*, 2005). We used GSEA with the Gene Ontology (GO) cellular component gene set. We found that particularly in valve endocardial cells and atrioventricular cardiomyocytes, endosomal and lysosomal pathways were activated in mutants compared to siblings (Fig. 6C). This suggests a coupling mechanism for the lack of functional lysosomes and relates to the increased number of autophagic vesicles found in the mutant atrioventricular canal (Fig. 3B and C). In this regard, transcriptional responses to activate autophagy and lysosomal biogenesis have been described both upon lysosomal damage (Bonam *et al*, 2019) and in the context of epithelial development (Tognon *et al*, 2016).

We also performed over representation analysis on the differentially expressed genes using gene ontology cellular components (Fig. S5A, Table S5). This analysis identifies pathways, which have an overrepresented number of differentially expressed genes (Wu *et al*, 2021). We found an enrichment of genes related to the vacuolar proton-transporting V-type ATPase complex in the mutant EnVCs (Fig. S5A, Table S5). This is consistent with the described balanced interaction between Spns1 and the lysosomal V−type ATPase (Sasaki *et al*, 2017), which maintains lysosome pH gradients by pumping protons into the lysosomal lumen in an ATP-dependent manner. Also, genes related to transmembrane transporter activity were overrepresented in EnVCs (Fig. S5A), and several membrane-related pathways were significantly activated in both EnVCs and atrioventricular CMs (Fig. 6D and E). These results suggest a role for *spns1* in membrane organization and go in line with its more recently described function as transmembrane phospholipid-transporter (He *et al*, 2022; Scharenberg *et al*, 2023). Interestingly, no enrichment or overrepresentation was found in proliferation or apoptosis-related signalling pathways when comparing the mutant and sibling transcriptome, suggesting that the observed phenotype is not due to a general cardiac development impairment, but is rather the result of specific alterations in morphogenesis.

During heart development in general, and valve formation in particular, a tight signalling interaction between myocardium and endocardium is required (Luxán *et al*, 2016; Qu *et al*, 2022). To understand the intercellular signalling governing the observed changes, we performed a ligand-receptor analysis using LIANA (LIgand-receptor ANalysis frAmework) pipeline on both the sibling and *nrs* mutant hearts (Fig. 6F, Table S6). LIANA compiles the ligand-receptor interaction analysis from various sources and provides an aggregate rank for each potential interaction (Dimitrov *et al*, 2022). We found that among the ten top-ranking ligand-receptor interactions involving cardiomyocytes and endocardial cells, four affected *notch1b*-signalling. Specifically, contactin 1a (*cntn1a*)-, *ybx1*-, *jag2*-and *adam10*-*notch1b* interactions were altered in mutant versus sibling cardiac cells, in cell populations near AV region (CM AVC, EnVCs).

A deeper analysis revealed that the cause of the altered *notch1b* interactome was a reduction in the proportion of *notch1b*-expressing endocardial cells as well as atrioventricular cardiomyocytes (Fig. S5B). These cells also revealed a reduction in net *notch1b* expression and in the cell number and expression levels for the notch ligand Jagb2, as well as altered expression of the Notch1-activating enzyme Adam10 in some endocardial subclusters (Fig. S5B). We also found that *alcama*-dependent interactions involving cardiomyocytes and endocardial cells were affected in the mutant (Fig. 5F), which could be explained in part by the reduced Alcam expression observed in the mutant AV endocardial cells (Fig. 4B).

### Endocardial specific overexpression of notch1a rescues heart development and function

Given the importance of *notch1b*-signaling to valve formation (Luxán *et al*, 2016; MacGrogan *et al*, 2018), and to validate our transcriptomics data, we decided to investigate the expression of *notch1b* in the *nrs* mutant using mRNA *in situ* hybridization. A high proportion of mutants revealed a mislocated or downregulated *notch1b* expression compared to siblings at both 2 and 3 dpf (Fig. 7A). Moreover, lineage tracing analysis using the *notch1*-activity fate mapping line *Tg(tp1:CreERt2)* revealed a lower number of *tp1*-derived^+^ cells in the mutant heart, confirming reduced *notch1* signalling (Fig. 7B). To functionally evaluate if the mutant phenotype was driven by *notch1* signalling, we decided to overexpress the constitutively active intracellular domain of Notch1 specifically in endocardial cells using a Gal4/UAS-mediated strategy and the transgenic line *Tg(UAS:myc-Notch1a-intra)*) (Scheer *et al*, 2002). We obtained single and double transgenic mutant larvae through the crossing of heterozygous single transgenic parents (Fig. 7C, *spns1^+/-^, Tg(fli1a:Gal4)* and *spns1^+/-^, Tg(UAS:notch1-intra*)) and compared the larval heart morphology at 4 dpf. Our results showed that expression of the active Notch1 receptor (Scheer *et al*, 2001) in the endocardium led to a significant increase in the ventricle area (Fig. 7D). Moreover, the overexpression of *notch1* reinstated to some extent the morphological and functional characteristics of the atrioventricular canal (Fig. 7E-G, Video S10), which contributed to a higher percentage of mutant larvae with normal blood flow (Fig. 7H). Interestingly, the rescue appeared to be specific for the AV valves, as observed in the degree of elongation of the OFT and AV valves in rescued larvae (Fig. 6F and G). Our results hence show that expression of *notch1* in the endocardium could partially rescue the myocardial *nrs* phenotype.

**Figure 7.**
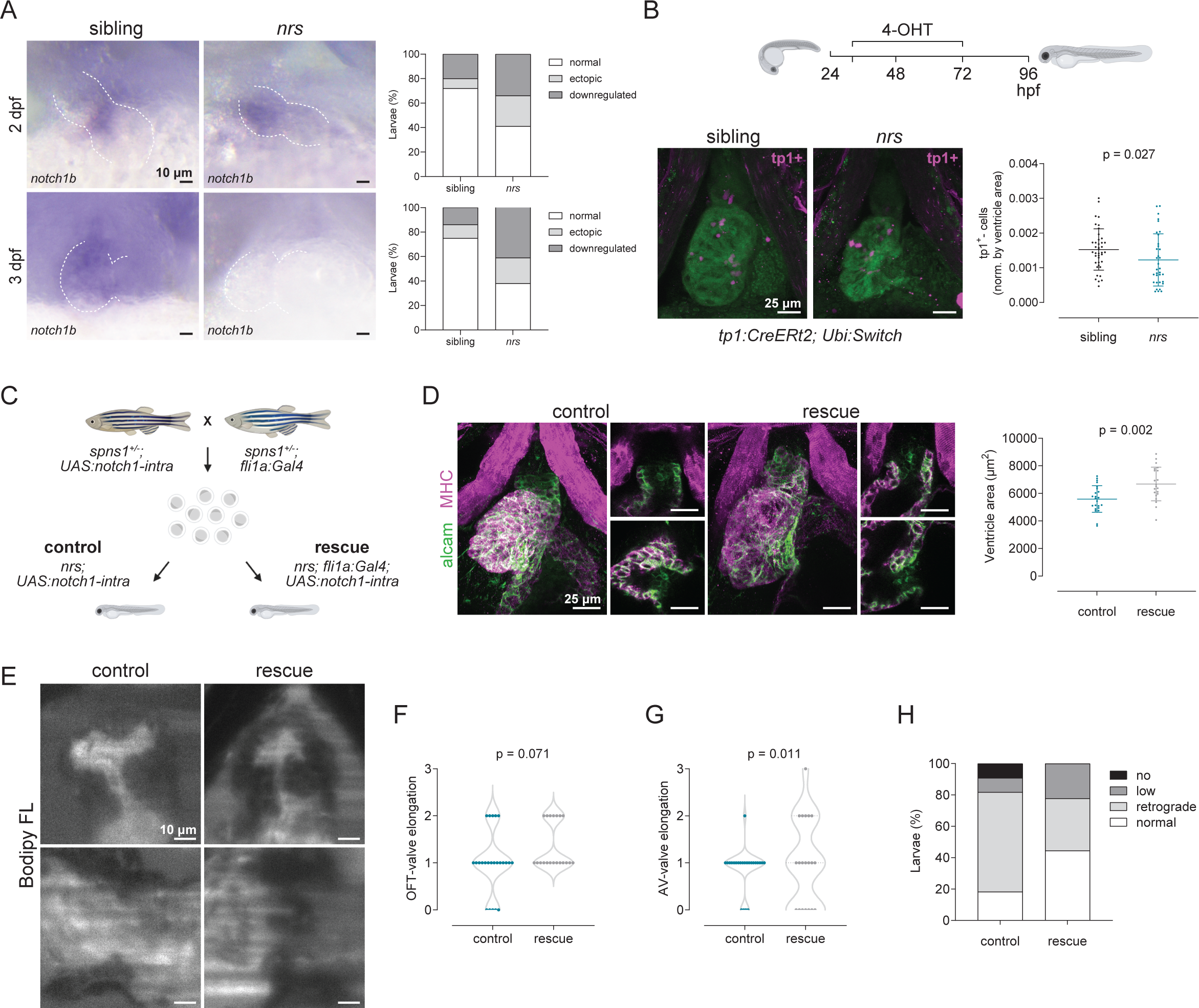
Endothelial specific overexpression of *notch1* rescues heart development and function. (**A**) The expression pattern of *notch1b* was assessed using mRNA *in situ* hybridization. A high proportion of *nrs* mutants show ectopic expression in the ventricle at 2 dpf, which differed from the localized atrioventricular expression in siblings. At 3 dpf, *notch1b* was downregulated in about 40% of mutant larvae. (**B**) Lineage tracing analysis using the tamoxifen (4-OHT) inducible *notch1*-activity fate mapping line *Tg(tp1:CreERt2)* revealed a lower number of *tp1*^+^-derived cells in the mutant heart, confirming reduced *notch1* signalling. Stack projections of whole mount immunostained hearts imaged by confocal microscopy are shown. Each dot represents one larva. Statistical analysis was performed using Mann-Whitney-U test. (**C**) Endothelial-cell specific overexpression of a constitutively active form of the Notch1 receptor in *nrs* larvae was achieved by outcrossing heterozygous single transgenic parents (*spns1^+/-^, Tg(fli1a:Gal4)* and *spns1^+/-^, Tg(UAS:notch1-intra)*). Rescue of the cardiac phenotypes observed in the mutant hearts was evaluated between control (*nrs, Tg(UAS:notch1-intra)*) and rescue (*nrs, Tg(fli1a:Gal4, UAS:notch1-intra)* conditions. (**D**) Whole-mount immunostaining of control and rescued *nrs* hearts showed that *notch1* overexpression significantly increased the ventricle size of the mutants (MHC: Myosin heavy chain). Each dot represents one larva. Statistical analysis was performed using Welch’s t-test. (**E-H**) Cardiac and valve function was evaluated in BODIPY™ FL C5-stained larvae using light-sheet microscopy. Qualitative parameters were considered to evaluate (**F**) OFT-and (**G**) AV-valve elongation and function (1: valve cushions, 2: delaminated valve cells, 3: elongated clapping valves). Each dot represents one valve. Chi-square test was applied to analyse differences between experimental groups. (**H**) Percentage of larvae presenting normal, low, retrograde or no blood flow.

## Discussion

Despite the clinical relevance of the pathophysiological connection between cellular homeostatic processes and congenital cardiac diseases, the role of autophagy-lysosomal degradation during cardiac development is understudied, partly because of limited options to visualize these dynamic processes *in vivo* and during the formation of specific cardiac structures, such as the heart valves. Furthermore, assessment of autophagic activity in the zebrafish has until now relied on the use of ectopic *lc3b* expression under the control of tissue-specific or semi-ubiquitous promoters (Lee *et al*, 2014; Mathai *et al*, 2017; Zhang *et al*, 2022; Chávez *et al*, 2020), which fails to recapitulate endogenous expression and expression in cell types that do not activate the used promoter.

In this work, we provide a detailed characterization of autophagosome and lysosome formation/accumulation during heart and valve development. The *Tg(mRFP-Lc3)* knock-in line established here now allows to assess spatiotemporal autophagosome formation at subcellular levels in zebrafish organ development, homeostasis and regeneration. In addition to confirming a sustained increase in autophagosome/autolysosome numbers in the myocardium between 48 and 96 hpf (Zhang *et al*, 2022), we observed a marked accumulation of autophagosomes/autolysosomes in the bulboventricular and atrioventricular endocardium at the developmental stage when cardiac valve formation begins. Furthermore, we uncovered a significant accumulation of *lamp2^+^* lysosomes and LysoTracker™ acidic vesicles in endocardial cells of the OFT and AVC, and their respective valves, as well as in AVC cardiomyocytes when valve delamination and elongation take place (72-96 hpf). Unfortunately, we were not able to evaluate lysosomal acidification using pH-sensitive dyes (Sasaki *et al*, 2014, 2017) with our imaging settings, which would have provided additional information regarding lysosomal function. Cardiac valve development is largely mediated by the transduction of mechanical forces, such as hemodynamic shear stress and reversing flow, to which endocardial cells in the AVC and OFT are exposed once the heart starts contracting (Steed *et al*, 2016a; Andrés-Delgado & Mercader, 2016). Interestingly, both oscillatory shear stress and disturbed flow induce autophagy in atherosclerotic lesions (Kheloufi *et al*, 2018) in a cytoprotective process that was recently described to implicate a lysosome associated protein (Canham *et al*, 2023). In line with this, our results indicate that a localized activation of autophagic processes supports the formation of the cardiac valves.

The *nrs* mutant model allowed us to investigate how impaired lysosomal function affects valve formation and function. The smaller and rounder ventricles described here for the *nrs*-mutant have also been observed upon *lamp2* loss of function (Dvornikov *et al*, 2019) and in a zebrafish knock-down for iduronate-2-sulfatase resembling mucopolysaccharidosis type II (Costa *et al*, 2017). Moreover, we found a severe impairment of both OFT and AVC valve development in *nrs* hearts, which affected the delamination and elongation of valves, and consequently their proper function. This correlated with the accumulation of autolysosomal contents observed in these cardiac regions in *nrs* mutants (Fig. 3). As to this, abnormal valve phenotypes have been associated with inherited lysosomal storage diseases, which are caused by genetic defects in lysosomal enzymes leading to a progressive lysosomal accumulation of substrates (Mueller *et al*, 2013; Ruiz-Guerrero & Barriales-Villa, 2018). The similarity of the phenotypes observed in *nrs* larvae and other human lysosomal disease zebrafish models (Costa *et al*, 2017; Yanagisawa *et al*, 2017), suggest that mutations in the *spns1* gene could lead to a similar pathology in humans. Importantly, our endocardial-specific rescue of the *nrs* valve phenotype shows, that *spns1*-dependet lysosomal function is required cell-autonomously in the endocardium for proper valve formation.

Endocardial cell volume decrease is required during zebrafish cardiac valve formation (Vignes *et al*, 2022), and indeed, our transcriptome analysis of the mutant heart revealed significant deviations in plasma membrane components (Fig. 6D and E). It would be plausible to propose that autophagic-degradation participates in the reorganization of cellular contents and plasma membrane remodelling during cardiac valve development. Two studies have recently identified Spns1 as a proton-dependent lysophosphatidylcholine transporter that is required for membrane phospholipid salvage and recycling (He *et al*, 2022; Scharenberg *et al*, 2023). These studies have shown that, similar to what we observed in *nrs* mutants, Spns1 deficiency in knockdown murine and zebrafish models, leads to the lysosomal accumulation of lipid molecules and consequently to overall lysosomal dysfunction due to luminal alkalization (He *et al*, 2022; Scharenberg *et al*, 2023). Altogether, these studies and the results presented in this work suggest a possible role for lysosomes in phospholipid metabolism and plasma membrane remodelling, directly affecting cardiac valve development.

Notch1b-signalling mediates early endocardial tissue patterning in the nascent valve cushions, as well as proliferation and remodelling during the later stages of valve development (MacGrogan *et al*, 2018). The ligand-receptor analysis and the experimental validation we performed show that lysosomal impairment affects *notch1b*-mediated communication within cardiomyocytes and endocardial cells, whereas endocardial-specific *notch1* overexpression rescues ventricle size and atrioventricular valve elongation in *nrs* mutants. Studies on stem cell expansion have described autophagy to modulate Notch signalling by regulating Notch1 degradation (Wu *et al*, 2016), whereas in neural stem cell asymmetrical divisions, differences in Notch signal activity depend on lysosome inheritance and the acidifying endolysosomal environment that allows the activating proteolytic cleavage of Notch1 (Bohl *et al*, 2022). Consistent with this, we find that impaired autophagosome processing and lysosome acidification in *nrs* reduces *notch1b* expression as well as *notch1*-signalling. Our results hence support previous findings showing that disruption of the lysosomal function affects Notch1-signaling (Lu *et al*, 2020; Tognon *et al*, 2016) and provides functional support for Notch1 being a central signalling pathway affected by aberrant endocardial lysosomal function during valve formation.

Finally, our results also show that endocardial cell organization is severely affected in *spns1* mutants. Autophagy and lysosomal processes are known to contribute to cell organization in other tissues (Saera-Vila *et al*, 2016; Vion *et al*, 2017; Moss *et al*, 2021) and to be tightly intertwined with ECM deposition and remodelling (Kim *et al*, 2012; Kawano *et al*, 2017; Schaefer & Dikic, 2021). For example, in the vasculature, where flow-induced endothelial cell alignment is disturbed by alterations in autophagy (Vion *et al*, 2017), or the cartilage, where autophagy impairments disorganize differentiating chondrocytes and their surrounding ECM (Moss *et al*, 2021). Indeed, lysosomal impairment in *nrs* mutants directly affected *alcama*-signalling, a cell-adhesion molecule for which several functional roles related to cell clustering, proliferation and migration have been described in developing tissues (Swart, 2002; Choudhry *et al*, 2011). Since proper valve function greatly depends on correct valve morphogenesis and structural integrity, which in turn is linked to valve pathologies (Hinton & Yutzey, 2011), it will be clinically relevant to understand whether autophagy-lysosomal processes play a structural role in the formation of functional mature valve tissue and investigate their potential therapeutic targeting.

## Materials and Methods

### Zebrafish husbandry

Experiments were conducted with zebrafish (*Danio rerio*) embryos and larvae between 48-and 96 hpf. Eggs were obtained by natural spawning using slope breeding tanks from adult zebrafish aged 4-18 months. Larvae were grown at 28°C in E3 medium containing 1-phenyl-2-thiourea (0.003%, PTU, Sigma-Aldrich, P7629-10,) to avoid pigmentation. Adult zebrafish were maintained at a maximal fish density of 5 per Liter and under the following environmental conditions: 27.5-28°C, 650-700 μs/cm, pH 7.5 and daily 10% water exchange. All experimental protocols were performed at the Institute of Anatomy (national license number 35 and license for the generation of genetically modified animals number G BE8/19). An overview of all transgenic lines used in this study with information about their origin can be found in Table 1. Transgenic zebrafish embryos and larvae were selected under a fluorescent stereoscope (Nikon, SMZ800N). Mutant *nrs* larvae were identified by their described opaque yolk phenotype (Sasaki *et al*, 2017) under a brightfield stereoscope. Genotyping was performed to confirm the mutant genotype in larvae, to select heterozygous *nrs* adult zebrafish and to identify Gal4^+^-transgenic zebrafish. The respective primer sequences can be found under Table S7.

**Table 1.**
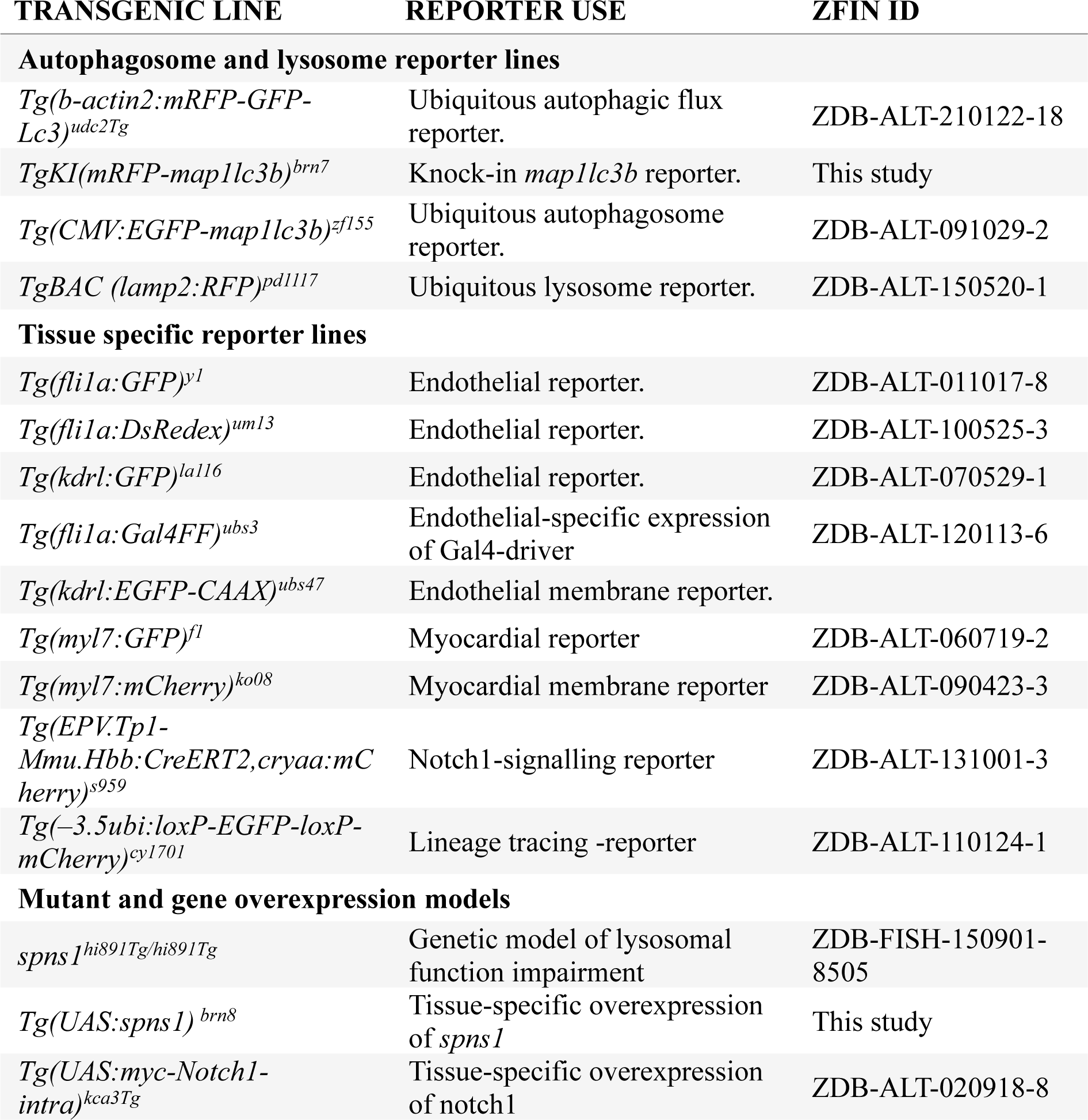
Transgenic zebrafish models used in the study.

### Generation of new transgenic lines

The endogenous autophagosome reporter line *TgKI(mRFP-lc3)* was generated following a previously described strategy for Cas9 protein-mediated oligonucleotide insertion (Gagnon *et al*, 2014). Briefly, gRNA candidates were designed based on a previously sequenced 432 bp region covering the first exon and intron of map1lc3b using online resources (http://crispor.tefor.net, https://eu.idtdna.com/site/order/designtool/index/CRISPR_CUSTOM). For gRNA selection, cutting efficiency was determined by T7-endonuclease assay (T7 Endonuclease I, New England Biolabs, M0302) according to the manufacturer’s protocol. Sequences for two ∼1 kb homology arms flanking the CRISPR targeting site were obtained by PCR (Q5® High-Fidelity DNA Polymerase, New England Biolabs, M0491) from parental zebrafish (AB strain) genomic DNA according to the manufacturer’s protocol. Three mutations were introduced to avoid CRISPR/Cas targeting through PCR and site-directed mutagenesis. The coding sequence of the fluorescent protein mRFP was obtained by PCR using the plasmid *pTol2-bactin2:mRFP-GFP-Lc3* (Chávez *et al*, 2020) as template. It was then modified to contain a short linker protein sequence (IDELNS) based on a similar construct (Kaizuka *et al*, 2016), to guarantee proper protein folding. Fragments were assembled into plasmid *pKHR4* (Hoshijima *et al*, 2016) (Addgene, 74592, Grunwald lab) by Gibson cloning to obtain the final repair template. To establish the knock-in line, fragment containing the repair template was obtained by enzymatic digestion with BamHI, HindIII and ScaI (FastDigest Restriction Enzymes, Thermofisher Scientific, FD0054, FD0504, FD0434), purified (DNA Clean & Concentrator, Zymo Research) and injected (25 ng/µl) along with Cas9 protein (0.3 mg/ml), gRNA (5.6 µM crRNA and tracerRNA, synthesized by IDT Integrated DNA Technologies) and 0.2 µM KCl (Sigma, P9333-500G) into one-cell stage embryos. The respective primers and gRNA sequences are listed in Table S7. Correct in-frame insertion of the *mRFP*-sequence into the zebrafish genome and expression of the endogenous fluorescently labelled Lc3 protein was confirmed by sequencing of gDNA and cDNA in F1-larvae, respectively. A schematic illustration of the repair template and the sequence alignment of the integrated transgene can be found in Fig. S1A and B. The line has been deposited at ZFIN under the name *TgKI(mRFP-map1lc3b)^brn7^.* The construct *pTol2-cry:mCherry-UAS:spns1* used for the tissue-specific overexpression of the wild-type *spns1* gene was generated by Gateway-mediated cloning. Briefly, total RNA was extracted from wild type zebrafish larvae (AB strain) through TRIzol RNA-extraction (Invitrogen, 10296010) and cDNA synthesis was performed using iScript Reverse Transcription SuperMix (Bio-Rad, 1708841). The coding sequence of the zebrafish *Spinster 1* (*spns1*) gene was amplified from cDNA by Phusion PCR using specific primers (Table S7) and subsequently cloned into the *pENTR/D-TOPO vector* (Invitrogen, K240020) to generate the *pME-spns1 vector*. The cloned sequence was verified by Sanger sequencing (Eurofins genomics). Multisite Gateway LR recombination was performed using the Gateway LR Clonase II Enzyme mix (Invitrogen, 11791020) to recombine the generated pME-spns1 plasmid, the Tol2Kit plasmids #327 (*p5E-UAS*) and #302 (*p3E-polyA*) (Kwan *et al*, 2007) and the destination vector pDestTol2pA2CrymCherry (Addgene, 64023; provided by Joachim Berger & Peter Currie (Berger & Currie, 2013)) to assemble the final p*Tol2-cry:mCherry-UAS:spns1* vector. The transgenic line *Tg(UAS:spns1, cry:mCherry)* was generated through Tol2-mediated recombination (Kawakami, 2007) by injecting one-cell stage embryos with the plasmid (25 ng/µl), 40 ng/ml Tol2 mRNA and 0.2 µM KCl (Sigma, P9333-500G). Overexpression of the wild-type *spns1* gene was confirmed by SYBR-green using cDNA from pools of 7-15 single (*Tg(UAS:spns1)*)and double transgenic (*Tg(fli1a:Gal4; UAS:spns1*)) *nrs* mutant larvae, using *rps11* (ribosomal protein S11) as housekeeping gene. The line has been deposited at ZFIN under the name *Tg(UAS:spns1, cry:mCherry)^brn8^*.

### gRNA synthesis and generation of *spns1* crispants

Two guide RNA (gRNA) were designed using CRISPRscan (Moreno-Mateos *et al*, 2015) based on their predicted efficiency to target in *spns1*, proximal location to its first exon, and low predicted off-target effects (Table S7). crRNA and tracerRNA were synthesized by IDT Integrated DNA Technologies, mixed in 1:1 ratio in Nuclease-Free Duplex Buffer and heated at 950C for 5 min to prepare the gRNA duplex solution. To generate *spns1* crispants, one-cell stage embryos obtained from crosses of heterozygous *nrs* mutant and wild type zebrafish were microinjected with a solution containing both gRNAs (4.5 µM each), Cas9 protein (0.3 mg/ml), P9333-500G) and 0.2 µM KCl (Sigma, P9333-500G). An injection solution without Cas9 protein was used as control. Cutting efficiency was verified via T7 endonuclease assay, and crispants were selected based on the opaque yolk phenotype.

### *In vivo* light sheet fluorescence microscopy

Embryos were raised until the desired developmental stage. For autophagosome and lysosome quantification, larvae were treated with 2 mM chloroquine to stop autophagic flux (3 h for autophagosome/lysosome quantification during development, 12 h to compare autophagosome and lysosome accumulation in *nrs* mutants). To label acidic vesicles/lysosomes, zebrafish were incubated for 3 h in 1 µM LysoTracker™ Deep Red (Invitrogen™, L12492) in E3 (DMSO). For the visualization of cardiac valves, larvae were incubated with 2 µM BODIPY™ FL C5-Ceramide (Thermo Fisher Scientific) in E3 (0.1% DMSO) overnight. Before imaging, larvae were washed 3 x 15 minutes with fresh E3 and anaesthetized using tricaine (0.08 mg/ml. pH7). Larvae were then embedded in 0.75% low-melt agarose with 0.04 mg/ml tricaine, transferred into a FEB-capillary and mounted inside a Leica-imaging chamber filled with E3-medium with tricaine. Imaging was performed with a Leica TCS SP8 digital light sheet (DLS) microscope equipped with a Hamamatsu Flash 4.0 V3 camera with 4.2 mpx using the following settings: 25X detection objective with NA 0.95 water immersion, 2.5X illumination objective, 5 mm illumination mirror. Images were obtained following XYTCZ acquisition mode with 1.14 ms exposure time to allow later retrospective gating.

### Retrospective gating, image processing and quantification

Images were processed using a HP-Zseries workstation (Dual Intel Xeone5-2667 v4 3.2 GHz, 256 GB, NVIDIA GeForce GTX 1080 Ti). Images were first exported from the lif-file into XYTC-tif files using the in house created macro *Export6D*. Retrospective gating was performed as previously described using the MATLAB (R2017a) tool BeatSync V2.1. to re-synchronize the 3D imaged hearts (3D-t reconstruction) (Marques *et al*, 2022). To improve the signal-to-noise ratio for quantification and visualization purposes, images were denoised using *Aydin* (https://pypi.org/project/aydin/) and the denoising algorithms *noise2self* and *butterworth*. Maximum intensity projections of the 3D image-stacks were obtained using ImageJ (Schneider *et al*, 2012) and images were cropped to the heart area using the in house created ImageJ macro *ClearOutside_3DnC*. For the quantitative analysis of autophagosomes and lysosomes in the developing heart, manual contour tracing and selection of regions of interest (ROIs; V: ventricle, OFT: outflow tract, AVC: atrioventricular canal) were performed in a first step and independent curation of the segmentations was performed in a second step (two-person control). Both segmentation and curation of image z-stacks (dimensions: xyzc, where c is the color channel of the respective endocardial, myocardial or ubiquitous tissue marker) were done in a slice-based approach using Napari and its labels layer functionality. The thickness of the traced contours varied between 1-2 (± 1) cell diameters, corresponding to the thickness of the specific tissue and ROI. To determine the median intensity of the respective vesicle reporter (autophagosomes, lysosomes) in each selected ROI, a custom Python script employing *skimage* algorithms was utilized for each imaged heart (N ≥ 5 for each transgenic line at each developmental time point). Relative differences of the median intensity ((AVC-V)/V, (OT-V)/V)) were estimated to compare autophagosomal/lysosomal vesicle accumulation in the AVC and OFT of the developing zebrafish heart in relation to the ventricle (V). All image processing scripts are available at “https://github.com/MercaderLabAnatomy/PUB_Chavez_et_al_2023”. To estimate autophagosome and lysosome accumulation in sibling and mutant hearts, the number of autophagosome and lysosome puncta were determined using the ImageJ *Particle analysis* plugin and normalized by the cross-sectional ventricle area. Similarly, estimation of cardiac morphological parameters (cross sectional ventricle area, circularity, outflow tract length and atrioventricular canal width) was performed using ImageJ. Semi-quantitative assessment of valve extension and heart rate at 4 dpf was done manually and blinded.

### Transmission electron microscopy

Zebrafish larvae were selected at 4 dpf, euthanized by tricaine-overdose and fixed with 2.5% glutaraldehyde (Agar Scientific, Stansted, Essex, UK) and 2% paraformaldehyde in 0.1 M Na-Cacodylate buffer (Merck, Darmstadt, Germany) with a pH of 7.44. Samples were fixed for at least 24 hours before being further processed. Samples were then washed with 0.1 M Na-Cacodylate buffer three times for 5 minutes each, postfixed with 1% OsO4 (Electron Microscopy Sciences, Hatfield, USA) in 0.1 M Na-cacodylate buffer at 4°C for 2 hours, and then washed in 0.1 M Na-Cacodylate buffer (Merck, Darmstadt, Germany) three times for 5 minutes each. Thereafter samples were dehydrated in 70%, 80%, and 96% ethanol (Grogg, Bern, Switzerland) for 15 minutes each at room temperature. Subsequently, samples were immersed in 100% ethanol (Merck, Darmstadt, Germany) three times for 10 minutes each, in acetone (Merck, Darmstadt, Germany) for two times for 10 minutes each, and finally in acetone-Epon (1:1) overnight at room temperature. The next day, samples were embedded in Epon (Sigma-Aldrich, Buchs, Switzerland) and left to harden at 60°C for 5 days. Sections were produced with an ultramicrotome UC6 (Leica Microsystems, Vienna, Austria). First, sequential semithin sections (1 um) were obtained and stained with a solution of 0.5% toluidine blue O (Merck, Darmstadt, Germany) for light microscopy to select cardiac regions comprising the atrioventricular canal and outflow tract. Upon selection, ultrathin sections (75 nm) were produced for electron microscopy, mounted on single slot copper grids, and stained with uranyless (Electron Microscopy Sciences, Hatfield, USA) and lead citrate (Leica Microsystems, Vienna, Austria) with an ultrostainer (Leica Microsystems, Vienna, Austria). Sections were examined with a transmission electron microscope (FEI Tecnai Spirit, Thermo Fisher Scientific).

### Immunofluorescence

For whole-mount immunofluorescence, zebrafish larvae were selected at the desired developmental stage, euthanized by an overdose of tricaine and fixed overnight at 4°C in 4% paraformaldehyde (PFA, EMS, 15710). Larvae were washed three times for 15 minutes each with PBS, dehydrated following a methanol gradient (25%. 50%, 75% methanol in PBS, 15 minutes each) and stored in 100% methanol for at least one day at −20°C. Larvae were re-hydrated (75%, 50%, 25% methanol in PBS-TritonX100 (0.3%), 10 minutes each) and washed twice with PBS-TritonX100 (0.3%) for 10 minutes. The permeabilization procedure consisted of treatment with Proteinase K (10 µg/ml) for 20 minutes, followed by 20 minutes post-fixation with 4% PFA and incubation in ice-cold Ethanol-Aceton 2:1 for 7 minutes, with three PBS-TritonX100 (0.3%) 10 minutes washes in between. Blocking of non-specific binding sites was performed by overnight incubation in blocking solution (5% BSA, 5% goat serum, 1% DMSO, 1% TritonX100) at 4°C. To achieve a better antibody penetration, the pericardial cavity was punctured using a tungsten needle and the incubation with first and secondary antibodies was performed overnight at 4°C. Primary antibody dilutions were prepared in PBS containing 5% BSA, 5% Goat serum and 0.002% sodium azide as follows: anti-alcam (1:100, ZN-8, DSHB), anti-MHC (1:100, MF20, DSHB), anti-mCherry (1:300, NBP2-25157, Novus Biologicals), anti-GFP (1:300, GFP-1010, Aves), anti-BrdU (1:200, abcam, ab6326). Goat-derived secondary antibodies (all obtained from Invitrogen, Thermofisher Scientific) were used in 1:300 dilution in PBS containing 5% BSA, 5% Goat serum: anti-mouse IgG1 - Alexa Fluor® 488 (A-21121), anti-mouse IgG2b - Alexa Fluor® 568 (A-21131), anti-rabbit (H+L) Superclonal™ - Alexa Fluor® 647 (A27040),anti-Chicken IgY (H+L) - Alexa Fluor® 488 (A-11039), anti-rat IgG (H+L)-Alexa Fluor® 647 (A21247).

### Heart function analysis

Transgenic *Tg(fli1a:GFP)* sibling and mutant larvae were selected at 3 dpf, and transferred to a flat-bottomed 96-well plate containing pre-molded 1% agarose pockets used to retain larvae in a ventral position. Imaging was performed using an automated microscopy platform (Acquifer Imaging Machine, Acquifer Germany). In this two-step imaging process, brightfield overview images of whole larvae were obtained using a 2x objective, which were used for the automated detection of the head region, and later, the estimation of the larval body length. Then, series of brightfield and green fluorescence images of the head region focused on the heart are acquired using a 10x objective to obtain a time series of the beating heart. Images were processed into hyper-stacks using the Acquifer built-in Fiji-software, and analyzed using previously described (Marques *et al*, 2022) in-house developed scripts to determine larval body length, heart rate (number of beats estimated during image acquisition,), ejection fraction (calculated from the difference between diastolic and systolic volume relative to the diastolic size), cardiac output (difference between diastolic and systolic volume multiplied by heart rate) and rhythmicity (RMSSD, evaluated by the root mean square of the successive differences between heart beats, which reflects the beat-to-beat variance). All FIJI plug-ins and Python scripts (“Heartbeat”, “Cardiac function”, “RMSSD”, “Size”) are available at “https://github.com/MercaderLabAnatomy/PUB_Chavez_et_al_2023”.

### Single nuclei gene expression library construction and analysis

Sibling and mutant larval hearts were obtained at 3 dpf by manual dissection using forceps and a tungsten needle as previously described (Singleman & Holtzman, 2011). Pools of approx. 50 hearts were collected within one hour in ice-cold Leibovitz’s L-15 Medium (Thermo Fisher Scientific) supplemented with 10% foetal bovine serum (Sigma-Aldrich, F7524), centrifuged for 4 minutes at 300 g, snap frozen and preserved in liquid nitrogen. Two replicates, each consisting of four pools, were obtained for each experimental group (sibling, mutant). Single nuclei suspensions containing 3800-4000 nuclei/ µL were prepared using the Chromium Nuclei Isolation Kit with RNase Inhibitor (10 x Genomics, PN-1000494) following the samples Prep User Guide (10 x Genomics, CG000505, Rev A). The Transposition, GEM generation & barcoding, reverse transcription, and preparation of the gene expression and ATAC libraries was performed according to the 4Chromium Next GEM Single Cell Multiome ATAC + Gene Expression User Guide (10x Genomics, CG000338 Rev F) with all stipulated 10x Genomics reagents. Nuclei suspensions were incubated in a Transposition Mix that includes a Transposase. At the end of the transposition step, GEM generation and barcoding was immediately performed, and a quenching reagent was added to each sample to stop the reaction. The reactions were then stored at −80° C. Samples were retrieved from storage, cleaned-up as stipulated in step 3.0 of the user guide. Thereafter, a pre-amplification steps was performed with 6 PCR cycles, followed by cDNA amplification and 3’ gene expression library workflow using 16 PCR cycles. Generated cDNA and both types of libraries were evaluated for quantity and quality using a Thermo Fisher Scientific Qubit 4.0 fluorometer with the Qubit dsDNA HS Assay Kit (Thermo Fisher Scientific, Q32851) and an Advanced Analytical Fragment Analyzer System using a Fragment Analyzer NGS Fragment Kit (Agilent, DNF-473), respectively. The cDNA libraries were pooled and sequenced with a loading concentration of 300 pM, asymmetric paired-end and dual indexed, using an illumina NovaSeq 6000 S1 Reagent Kit v1.5 100 cycles (illumina, 20028319) on an illumina NovaSeq 6000. The read set-up was as follows: read 1: 29 cycles, i7 index: 10 cycles, i5: 10 cycles and read 2: 89-91 cycles. The quality of the sequencing runs was assessed using illumina Sequencing Analysis Viewer (illumina version 2.4.7) and all base call files were demultiplexed and converted into FASTQ files using illumina bcl2fastq conversion software v2.20. A minimum of 50’000 reads/cell were generated for each library. All steps were performed at the Next Generation Sequencing Platform, University of Bern. The raw sequencing data was processed using cellranger v6.0 (Zheng *et al*, 2017). For alignment, we used Danio rerio Genome assembly GRCz11 v109 from Ensembl (Cunningham *et al*, 2022). The counts were then processed in R (v4.0) using Seurat (v4.0) and tidyverse (Hao *et al*, 2021; Stuart *et al*, 2019; Wickham *et al*, 2019). Possible doublets were removed using SingleCellExperiment and scDblFinder (Amezquita *et al*, 2020; Germain *et al*, 2022). Cells with a minimum of 200 genes and less than 5000 genes per cells were chosen for quality purposes. The final number of cells used for further processing was as follows: sib_1: 1673 cells, sib_2: 2193 cells, mut_1: 1643 cells, mut_2: 1940 cells. All possible mitochondrial reads (due to possible contamination) were removed before downstream processing. The samples were integrated using the Seurat Feature Integration pipeline and checked for batch correction. For the identification of the various cell types, marker genes provided in several publications (Grimes & Kirby, 2009; Goddard *et al*, 2017; Duchemin *et al*, 2019; Burkhard & Bakkers, 2018; Fontana *et al*, 2020; Queen *et al*, 2023; Ma *et al*, 2021) and databases in EnrichR were used (Chen *et al*, 2013; Kuleshov *et al*, 2016; Xie *et al*, 2021). Differential expression analysis for each cell type was performed using pseudobulk, EdgeR-LRT with the Libra package (Squair *et al*, 2021). Differentially expressed genes were converted to mouse orthologous genes with biomaRt and then subjected to pathway enrichment using overrepresentation analyses and Gene Set Enrichment Analyses (GSEA) with the Gene Ontology terms with the help from ClusterProfiler package (Durinck *et al*, 2009; Wu *et al*, 2021; Yu *et al*, 2012). To identify the ligand receptor interactions, we used LIANA that consolidates multiple ligand receptor databases and provides an aggregate rank (Dimitrov *et al*, 2022). Graphs were plotted using Seurat, ggplot2, ggpattern, viridis and dittoSeq (Bunis *et al*, 2021; Stuart *et al*, 2019).

### Data availability

The raw data is deposited on GEO under GSE (GEO accession number: GSE246850, Reviewer’s access code: czqhogweblyfnyt). The analyzed data is made available under a shiny app made using ShinyCell with additional modifications (Ouyang *et al*, 2021). The code for analyses is available at “https://github.com/MercaderLabAnatomy/PUB_Chavez_et_al_2023”.

### Statistical analysis

Statistical analysis was carried out using GraphPad Prism7. Kruskal-Wallis test was applied to analyse statistical differences in autophagosome and lysosome quantifications. Welch’s t-test and Mann-Whitney-U test were applied to compare differences between sibling and mutant or control and rescue conditions upon assessment of normal distribution. The Chi-square probability distribution test was applied to analyse categorical variables (qualitative assessment of valve elongation). Qualitative assessment of valve morphology and function was performed prior to the determination of the larvae genotype to attain a blinded analysis. All snRNAseq transcriptome analyses were done using differential genes with p-values less than 0.05.

---

**Figure S1: Characterization of *TgKI(mRFP-Lc3)* knock-in line and supporting information for autophagosome formation and lysosome accumulation during heart development.**

(**A**) Schematic visualization of the construction of newly established mRFP-Lc3 endogenous autophagosome reporter line, showing the sequence alignment of the *map1lc3b* zebrafish orthologue, the constructed repair template and knock-in transgenic line confirming the insertion of the reporter transgene before Exon 1. A silent mutation in Exon 1 (T>C) and three mutations incorporated into the repair template (orange) to avoid its CRISPR/Cas targeting were inserted in the transgenic line. The sequence of the guide RNA and PAM (underlined) are shown. (**B**) Overview confocal microscopy image of a *TgKI(mRFP-Lc3)* larva at 4 dpf. (**C**) Schematic representation of the transcript of mRFP-tagged Lc3 and sequencing primers. The correct in-frame insertion of the transgene was confirmed by sanger sequencing and (**D**) by amplification from cDNA of F1 larvae. (**E**) Quantification of the median fluorescence intensity of mRFP^+^-autophagosomes/autolysosomes in specific cardiac tissues at different developmental stages using ubiquitous (*Tg(b-actin2:mRFP-GFP-Lc3)*), endocardial (*Tg(fli1a:GFP, mRFP-Lc3)*) or myocardial (*Tg(myl7:GFP, mRFP-Lc3)*) reporter lines. Each point represents one larva (Kruskal-Wallis test, *p ≤ 0.05. **p ≤ 0.01). (**F**) The transgenic line *Tg(postnb:citrine)* was used to complement the description of autophagic processes in the developing heart and locate mRFP^+^-autophagosomes/autolysosomes in periostin-positive cells (*postnb^+^*) covering the outflow tract. (**G**) Quantification of the median fluorescence intensity of LysoTracker^TM^ DeepRed (ubiquitous) and mRFP-labelled lysosomes in the endocardium (*Tg(kdrl:EGFP-CAAX, mRFP-Lc3)* and myocardium (*Tg(myl7:GFP, mRFP-Lc3)*) at different developmental stages. Each point represents one larva (Kruskal-Wallis test, *p ≤ 0.05. **p ≤ 0.01). (**H**) Localization of LysoTracker^+^ lysosomes in periostin-positive cells (*postnb^+^*) in the outflow tract.

**Figure S2: *nrs* mutant heart presents morphological and functional abnormalities**

(A) Transmission electron microscopy of the sibling and *nrs* myocardium at 3 dpf. Overall, the distribution, size and shape of mitochondria (m) was not affected in mutant cardiomyocytes. Similarly, no evident differences in the organization or integrity of actin fibres (a) were observed between sibling and *nrs* heart. However, the accumulation of partially degraded lysosomal contents was observed within the myocardium, above all in atrioventricular cardiomyocytes. However, this was not as frequent or considerable as in the endocardium. (**B and C**) Confocal microscopy of immunostained larval hearts shows a significantly smaller and rounder (circularity index closer to 1) ventricle in *nrs* mutants at both 3 and 4 dpf. The measured ventricle cross sectional area was normalized to the average body size of sibling and mutant larvae to determine the organ specific phenotype. Each point represents one larva. Statistical analysis was performed using Mann-Whitney-U test. (**D-G**) Heart function was evaluated in sibling and mutant larvae at 3 dpf using a high-throughput imaging system by assessing the following parameters: (**D**) heart rate, (**E**) cardiac output, (**F**) ejection fraction and (**H**) degree of rhythmic beating (RMSSD). Each point represents one larva. (**D-G**) Statistical analysis was performed using Mann-Whitney-U test or Welch’s t-test depending on data distribution.

**Figure S3: *spns1* knock-down replicates cardiac phenotypes observed in *nrs* mutants**

(A) Brightfield images show the differences in the shape and appearance of the yolk in siblings, *nrs* mutants, control injected larvae (control) and *spns1* crispants (crispr). The *nrs* mutant phenotype is characterized by a round opaque yolk and regression of the yolk extension. (B) Confocal microscopy images of whole mount immunostained control and *spns1* crispant hearts. (**C and D**) Quantification of ventricle size and circularity index of control larvae and in *spns1* crispants at 4 dpf. Each point represents one larva. Statistical analysis was performed using Mann-Whitney-U test. (**E**) Valve development and morphology was evaluated in *Tg(fli1a:DsRed)* transgenic *spns1* crispant and control larvae at 4 dpf. The percentage of larvae with valve leaflets defined by alcam^+^-endothelial cells is shown. (**F-J**) Valve and heart function was evaluated in BODIPY™ FL C5-Ceramide stained *spns1* crispants and control injected larvae at 4 dpf. (**G and H**) A semi-quantitative assessment of valve morphology and movement revealed an impairment in valve elongation in *spns1* crispants. Each dot represents one valve. Chi-square test was applied to analyse differences between experimental groups. (**I**) Percentage of larvae presenting normal, low, retrograde or no blood flow. (**J**) Heart rate estimated by number of beats during a 150-frame acquisition. Each dot represents one larva. Statistical analysis was performed using Mann-Whitney-U test.

**Figure S4: Cell populations markers as identified in snRNAseq analysis**

(**A**) snRNAseq transcriptomic analysis of sibling and *nrs*-mutant hearts at 3 dpf allowed to identify several cardiac populations. (**B and C**) Dot plots depicting the cell markers that identify the various cell types as observed in snRNASeq for all the cell types and the subclusters of endocardial cells (**B**) and cardiomyocytes (**C**). Genes involved in valve development or genes which have been previously suggested as valve cell markers were used as reference to identify endocardial valve cells (EnVCs) and atrioventricular cardiomyocytes (CMs AVC) from within each respective endocardial and myocardial cluster.

**Figure S5. Overrepresentation analysis using Gene Ontology analysis for endocardial valve cells (EnVCs) and differential expression of *notch1b* signalling related genes**

(**A**) Overrepresentation analysis using gene ontology cellular components signals a significant effect in endosomal-lysosomal trafficking in EnVCs due to loss of *spns1.* (**B**) Bubble plot comparing the expression of *notch1b*-signalling related genes in atrioventricular cardiomyocytes and endocardial cell subclusters in sibling and mutant larvae.

**Video S1: Live-imaging of a *Tg(b-actin2:mRFP-GFP-Lc3)* zebrafish larval heart at 96 hpf.** The video shows the z-stack projection, one z-plane and a sequence of the stack images acquired by 3D+t light-sheet microscopy and upon reconstruction by retrospective gating. The cytoplasmic signal of GFP (green) as well as discrete mRFP^+^ autophagosomes/autolysosomes (magenta puncta) are sparsely distributed throughout the ventricle (V). Accumulation of both GFP/mRFP^+^ autophagosomes (white puncta) and mRFP^+^ autolysosomes can be distinguished the outflow track (OFT) and atrioventricular canal (AVC), suggesting increased autophagic activity in these regions (yellow arrows).

**Video S2: Live-imaging of a double transgenic *Tg(fli1a:GFP, mRFP-Lc3)* zebrafish larval heart at 96 hpf.** The knock-in endogenous autophagosome reporter *Tg(mRFP-Lc3)* allows to observe the accumulation (white arrows) of mRFP^+^ autophagosomes/autolysosomes (magenta puncta) in the endocardium (*Tg(fli1a:GFP,* yellow), particularly in endocardial valve cells of the outflow track (OFT) and atrioventricular canal (AVC). A z-stack projection, one z-plane and a sequence of the stack images acquired by 3D+t light-sheet microscopy and upon reconstruction by retrospective gating are shown (V: ventricle).

**Video S3: Live-imaging of a double transgenic *Tg(myl7:GFP, mRFP-Lc3)* zebrafish larval heart at 96 hpf.** The knock-in endogenous autophagosome reporter *Tg(mRFP-Lc3)* allows to observe the accumulation (white arrows) of mRFP^+^ autophagosomes/autolysosomes (magenta puncta) in cardiomyocytes (*Tg(myl7:GFP,* yellow) in the outflow track (OFT) just below the clapping valve, and at the base of the valves in the atrioventricular canal (AVC). The video shows a z-stack projection, one z-plane and a sequence of the stack images acquired by 3D+t light-sheet microscopy and upon reconstruction by retrospective gating. (V: ventricle).

**Video S4: Live-imaging of LysoTracker^TM^ DeepRed labelled lysosomes in a *Tg(cmv:EGFP-Lc3)* zebrafish heart at 96 hpf.** The autophagosome reporter *Tg(cmv:EGFP-Lc3)* shows high fluorescence (green) in the outflow tract (OFT) and the atrioventricular canal (AVC) suggesting greater accumulation of autophagic vesicles in these cardiac regions. Similarly, a significant accumulation of LysoTracker-labelled lysosomes (magenta puncta) is observed in the developing cardiac valves (yellow arrows). Altogether these results indicate a high rate of autophagic-lysosomal processing during valve development. The video shows a z-stack projection, one z-plane and a sequence of the stack images acquired by 3D+t light-sheet microscopy and upon reconstruction by retrospective gating. (V: ventricle).

**Video S5: Live-imaging of a double transgenic *Tg(kdrl:EGFP-CAAX, lamp2:RFP)* zebrafish larval heart at 96 hpf.** The video shows a z-stack projection, one z-plane and a sequence of the stack images acquired by 3D+t light-sheet microscopy and upon reconstruction by retrospective gating. Lysosomes (mRFP^+^ puncta) are observed throughout the larval heart (V: ventricle), yet they are most densely localized in *krdl^+^* endocardial cells and valve cells (white arrows) of the outflow tract (OFT) and atrioventricular canal (AVC). This hints towards special lysosomal function requirements for cells in these regions.

**Video S6: Live imaging of a double transgenic *Tg(myl7l:GFP, lamp2:RFP)* zebrafish larval heart at 96 hpf.** The video shows a z-stack projection, one z-plane and a sequence of the stack images acquired by 3D+t light-sheet microscopy and upon reconstruction by retrospective gating. Lysosomes (mRFP^+^ puncta) are observed to be distributed throughout the larval ventricle (V), yet they appear to be denser (white arrows) in myocardial cells (yellow) adjacent to the base of the valves in the outflow tract (OFT) and atrioventricular canal (AVC), suggesting higher lysosomal activity on these cardiac regions during the formation of their respective valves.

**Video S7: Live imaging of double transgenic *Tg(kdrl:GFP, myl7:mCherry)* sibling and *nrs* zebrafish hearts at 96 hpf.** The video shows the z-stack projections of beating sibling and *nrs* hearts to exemplify the constrained movement of the mutant endocardium (green) compared to the sibling. In contrast, the myocardium (magenta) contractility does not seem to be as affected in *nrs* mutants. Further, compared to the sibling heart, the outflow tract (white arrow) in the mutant heart shows limited capacity of expansion.

**Video S8: Live-imaging of BODIPY™ FL C5-Ceramide-stained sibling and *nrs* zebrafish hearts at 96 hpf.** The video allows to compare the development, morphology and movement of sibling and *nrs* mutant bulboventricular (OFT) and atrioventricular (AVC) cardiac valves. In contrast to the elongated, detached and clapping valves preventing retrograde blood flow in sibling hearts, mutant valves have hardly delaminated or elongated by 96 hpf and they cannot properly close.

**Video S09: Live-imaging of BODIPY™ FL C5-Ceramide-stained control (*nrs, Tg(UAS:spns1)*) and rescue (*nrs, Tg(fli1a:Gal4, UAS:spns1)* mutant zebrafish hearts at 96 hpf.** Endothelial-specific overexpression of the wild-type *spns1* gene partially rescues cardiac valve development and function. Above all, rescued atrioventricular (AVC) valves are able to properly open and close, and thus support unidirectional blood flow. (OFT: outflow tract).

**Video S10: Live-imaging of BODIPY™ FL C5-Ceramide-stained control (*nrs, Tg(UAS:notch1a-intra)*) and rescue (*nrs, Tg(fli1a:Gal4, UAS:notch1-intra)* mutant zebrafish hearts at 96 hpf.** Endothelial-specific overexpression of a constitutively active version of the Notch1 receptor promotes the delamination and elongation of the mutant atrioventricular (AVC) cardiac valves, and to a lesser degree the bulboventricular (OFT) valves. This supports the closure of the valves and prevents retrograde flow.

**Table S1:** Marker genes for cell populations.

**Table S2:** Valve-related genes expressed in endocardial and myocardial subpopulations.

**Table S3:** Differentially expressed genes of all cell clusters between *nrs* mutants and siblings.

**Table S4:** Gene set enrichment analysis using Gene Ontology Cellular Components for differential genes in each cluster/cell type. Missing cell types in the file indicate no enrichment could be obtained for that cell type.

**Table S5:** Over-representation Analysis using Gene Ontology Cellular Components for differential genes in each cell cluster/cell type. Missing cell types in the file indicate no enrichment could be obtained for that cell type.

**Table S6:** Ligand Receptor analysis using LIANA with scores for ligand-receptor pairs possible for each cluster/cell type.

**Table S7:** Primers and gRNA sequences used in the study.

## Acknowledgements

We thank M. Affolter and M. Bagnat for providing the *TgBAC (lamp2:RFP)pd1044* zebrafish line. Electron microscopy sample preparation and imaging were performed with devices provided by the Microscopy Imaging Center (MIC) of the University of Bern. Special thanks to B. Haenni for his support with sample processing. We thank Xavier Langa, Anna Gliwa and Ahmet Kürk for their assistance in zebrafish husbandry. Many thanks to Inês Marques for her experimental advice, and to Ayisha Marwa MP and Nick Kirschke for their support during sample collection. Illustrations and schemes were created with BioRender.com and with the support of S. Chávez Rosas.

## Conflict of Interests

The authors declare that they have no conflict of interest.

## Funding sources

This work was supported by grants 310030L_182575 from the Swiss National Science Foundation and H2020-SC1-2019-Single-Stage-RTD REANIMA-874764 to N.M., SELF2020-03 from University of Bern, Swiss Life research grant 2021 and ESC Basic Research Fellowship 2022 to M.N.C.

## Author contribution

**M.N.C.:** designed and performed the experiments, analyzed and interpreted results, wrote the manuscript, secured funding.

**P.A.:** performed snRNAseq bioinformatics analysis and generated related figures, gave statistics advice, contributed to writing the manuscript.

**A.E.:** contributed to optimization of imaging set-ups, heart function analysis and image processing pipelines.

**M.M.**: performed image segmentation, supported quantification analysis, and contributed to writing the manuscript.

**R.A.M.**: designed and generated the construct to establish the *Tg(UAS:spns1)* transgenic line and contributed to writing the manuscript.

**N.M.:** supervised the research, supported experimental design and the interpretation of the results, wrote the manuscript, secured funding.

## Notes

https://drive.google.com/drive/folders/1k7b5BNYztult-6XEmIRow25_Fzpg1UEF?usp=drive_link

### Competing Interest Statement

The authors have declared no competing interest.

