## Supplementary figures and images for "Spns1-dependent endocardial lysosomal function drives valve morphogenesis through Notch1-signaling"

### Supplementary_Figures

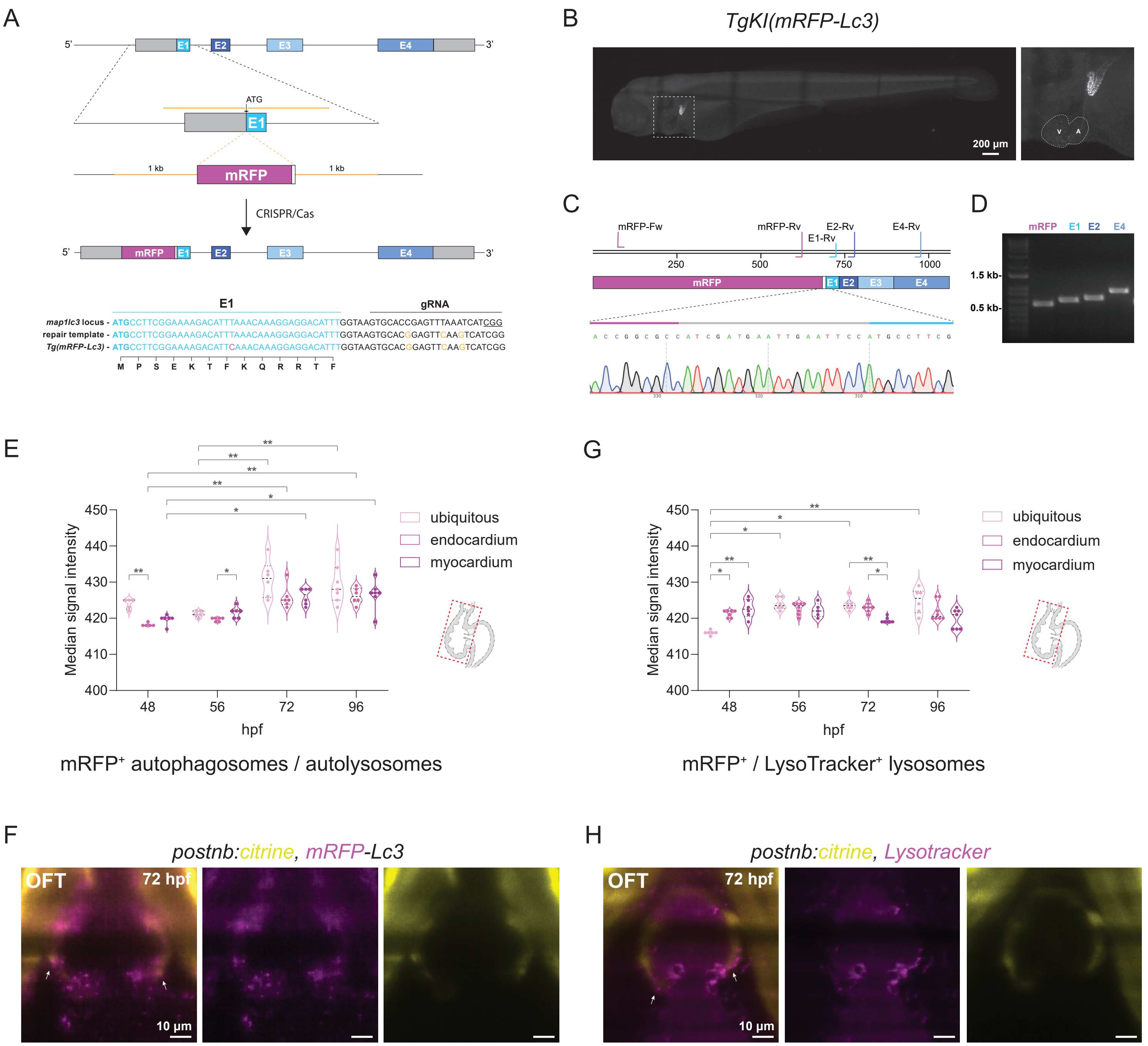

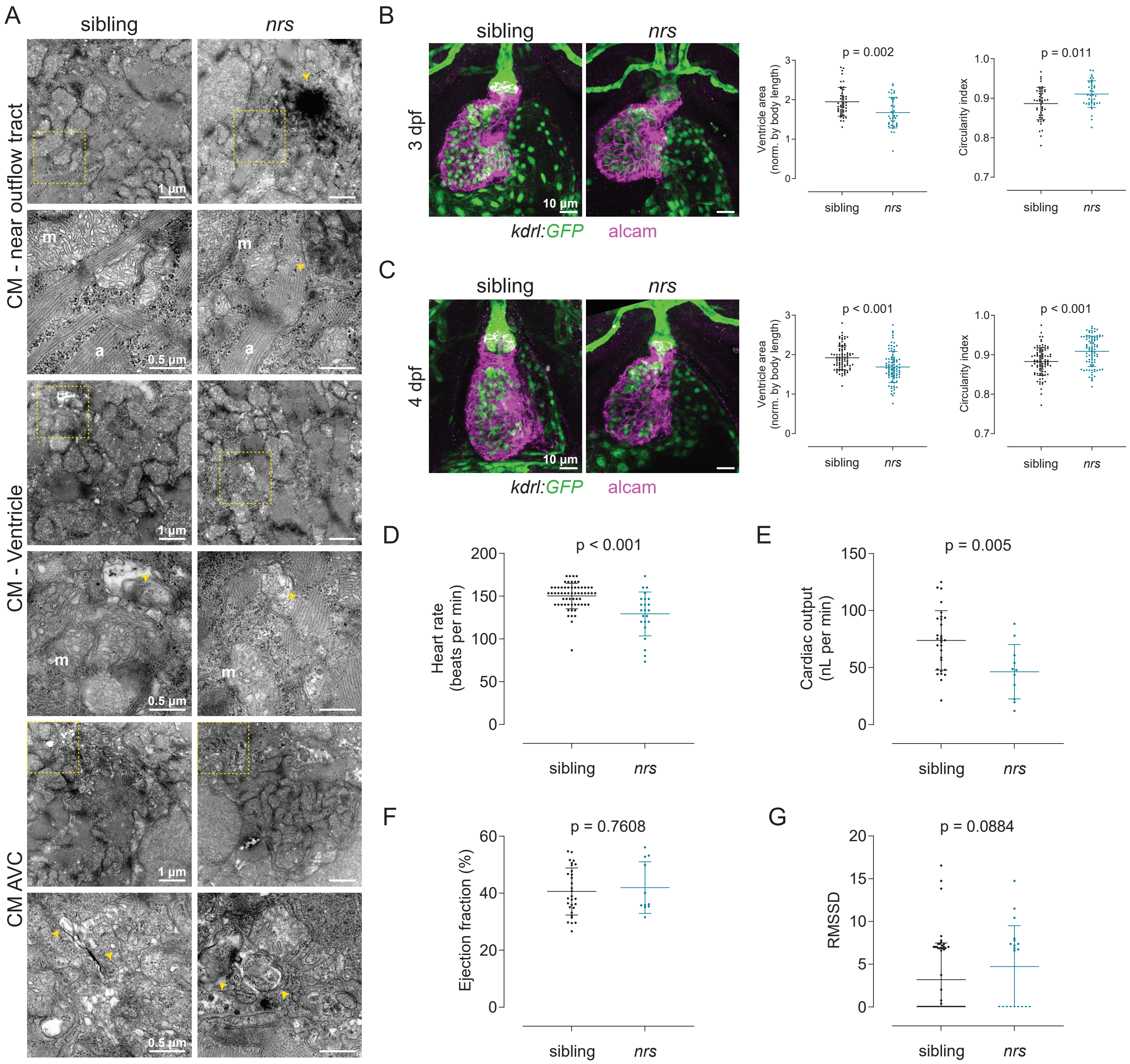

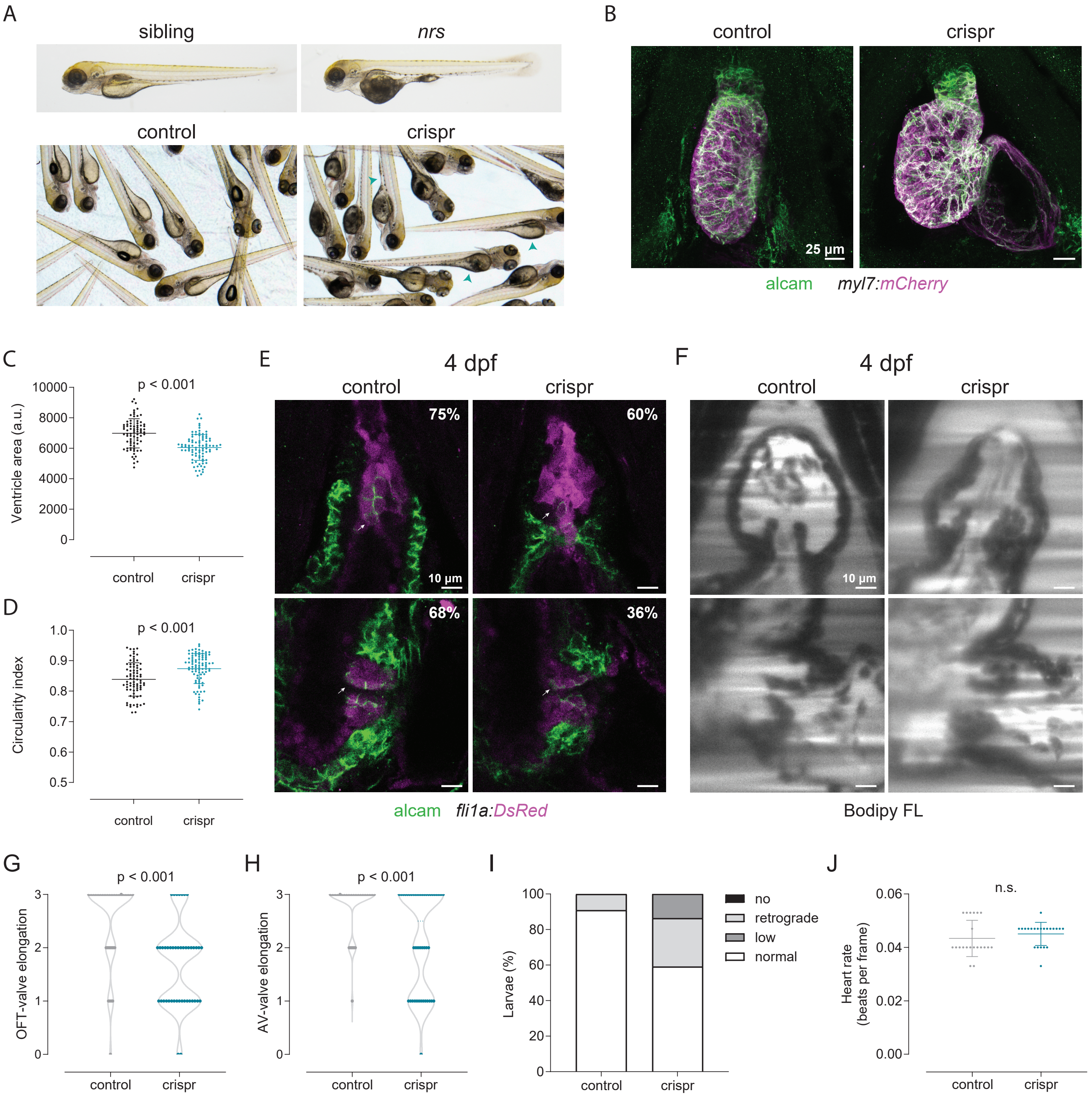

#### Supp. Fig 4

A

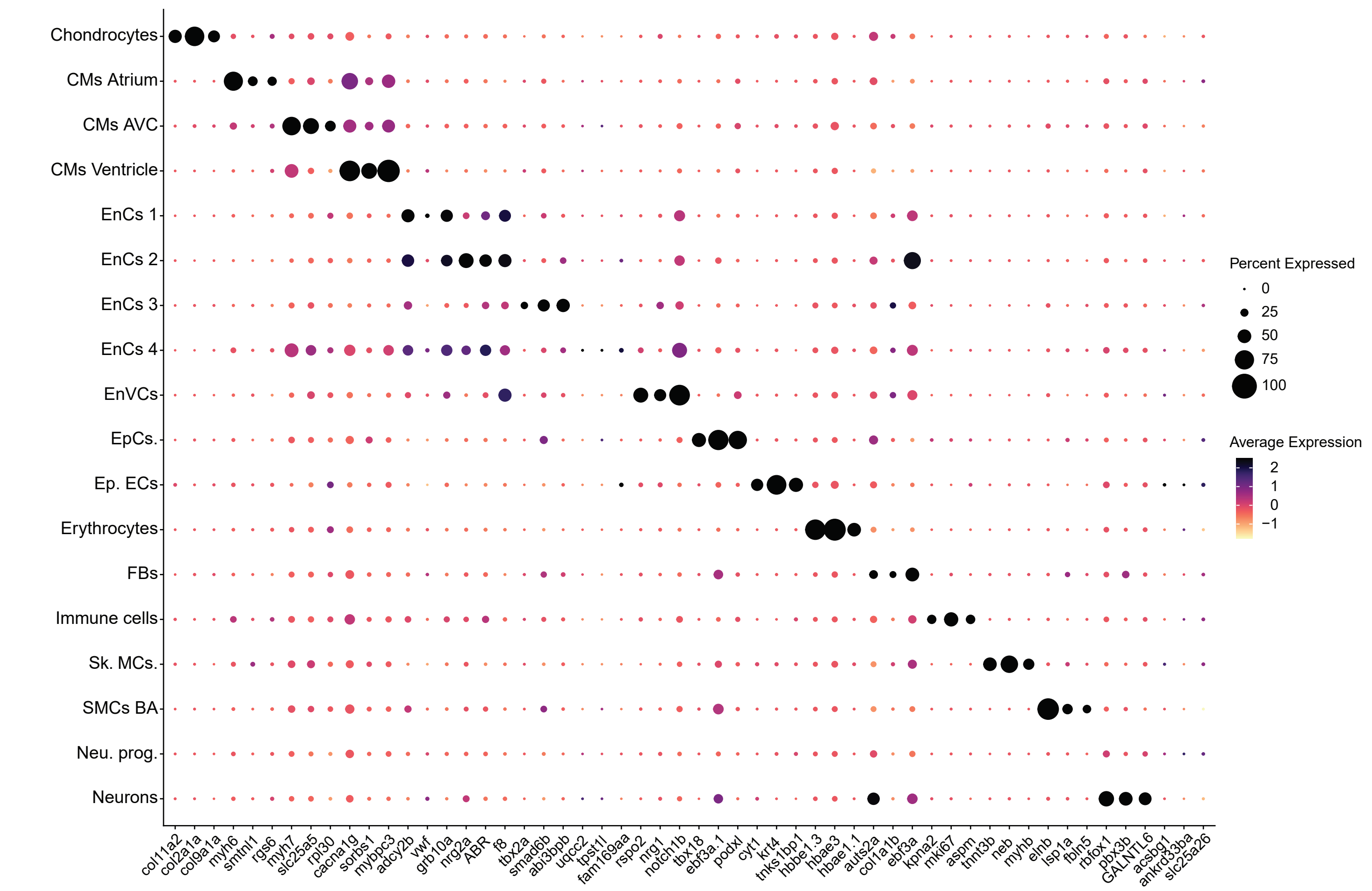


B

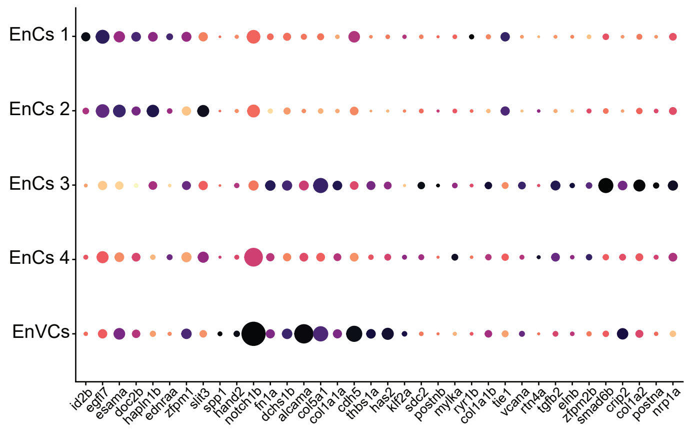


C

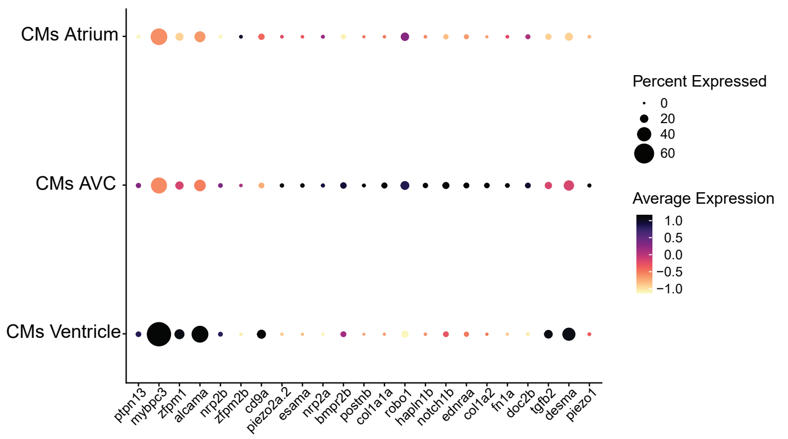

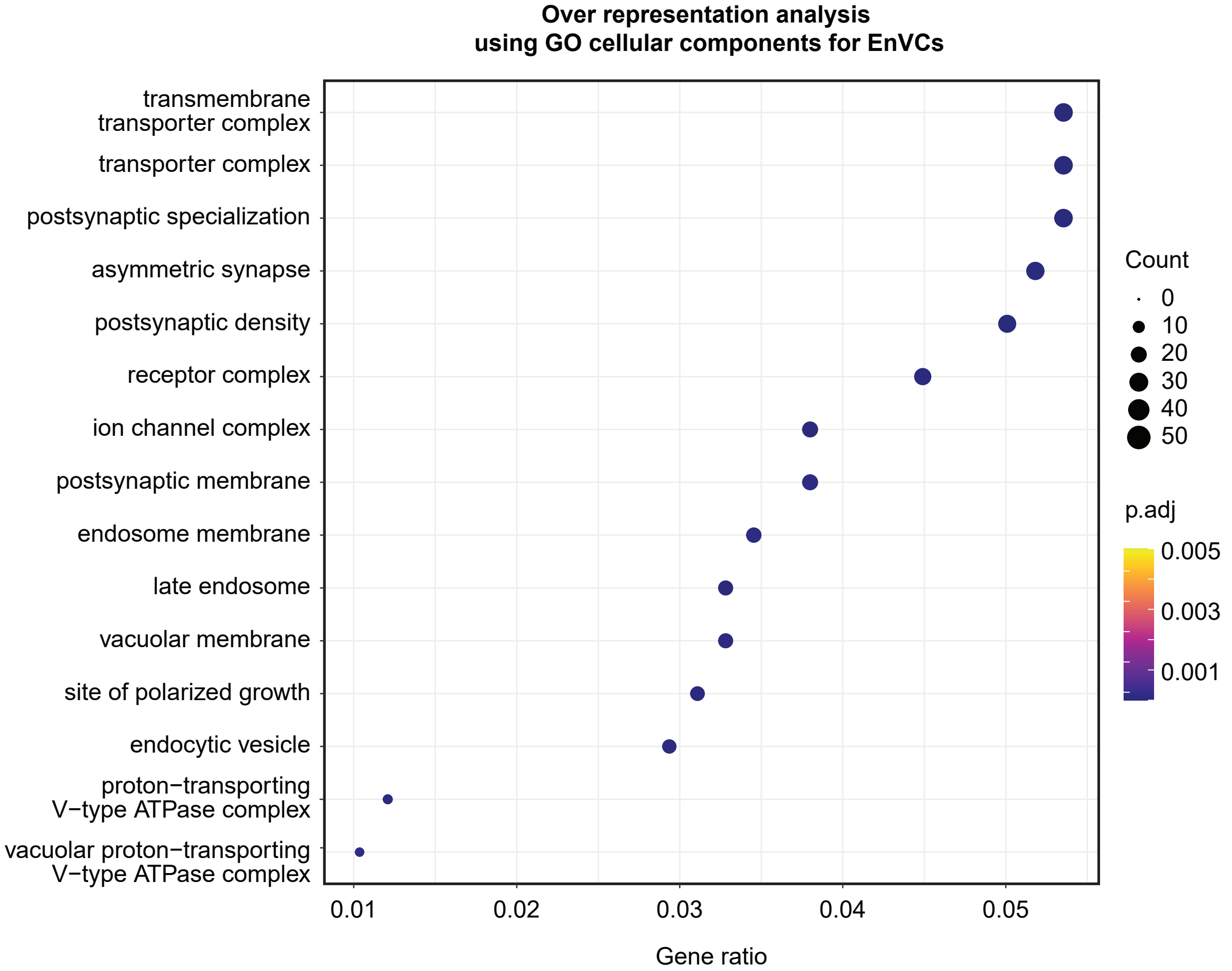

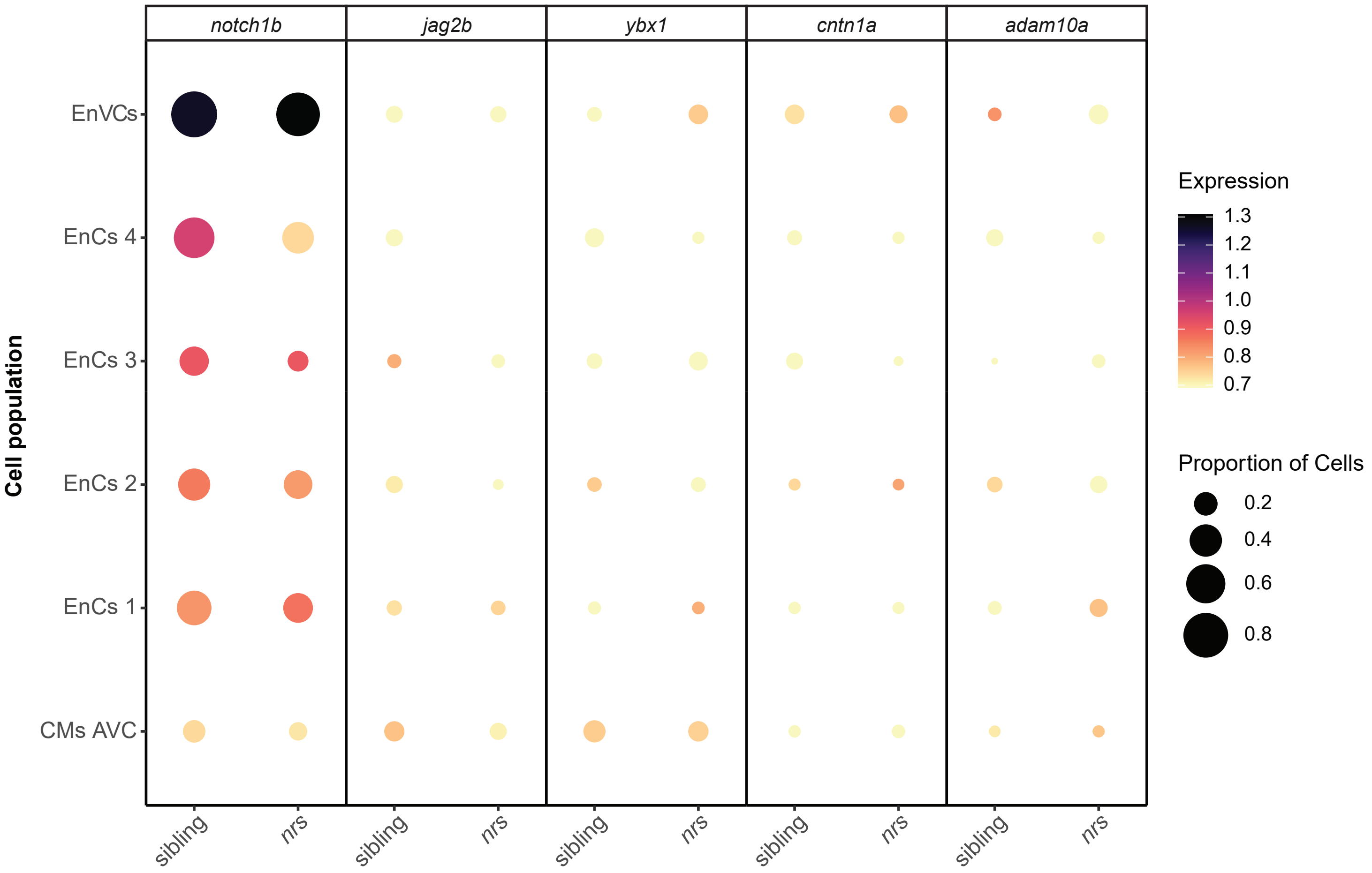
